# SMAUG: Analyzing single-molecule tracks with nonparametric Bayesian statistics

**DOI:** 10.1101/578567

**Authors:** J.D. Karslake, E.D. Donarski, S.A. Shelby, L.M. Demey, V.J. DiRita, S.L. Veatch, J.S. Biteen

## Abstract

Single-molecule fluorescence microscopy probes nanoscale, subcellular biology in real time. Existing methods for analyzing single-particle tracking data provide dynamical information, but can suffer from supervisory biases and high uncertainties. Here, we introduce a new approach to analyzing single-molecule trajectories: the <u>S</u>ingle-<u>M</u>olecule <u>A</u>nalysis by <u>U</u>nsupervised <u>G</u>ibbs sampling (SMAUG) algorithm, which uses nonparametric Bayesian statistics to uncover the whole range of information contained within a single-particle trajectory (SPT) dataset. Even in complex systems where multiple biological states lead to a number of observed mobility states, SMAUG provides the number of mobility states, the average diffusion coefficient of single molecules in that state, the fraction of single molecules in that state, the localization noise, and the probability of transitioning between two different states. In this paper, we provide the theoretical background for the SMAUG analysis and then we validate the method using realistic simulations of SPT datasets as well as experiments on a controlled *in vitro* system. Finally, we demonstrate SMAUG on real experimental systems in both prokaryotes and eukaryotes to measure the motions of the regulatory protein TcpP in *Vibrio cholerae* and the dynamics of the B-cell receptor antigen response pathway in lymphocytes. Overall, SMAUG provides a mathematically rigorous approach to measuring the real-time dynamics of molecular interactions in living cells.

**Statement of Significance:** Super-resolution microscopy allows researchers access to the motions of individual molecules inside living cells. However, due to experimental constraints and unknown interactions between molecules, rigorous conclusions cannot always be made from the resulting datasets when model fitting is used. SMAUG (Single-Molecule Analysis by Unsupervised Gibbs sampling) is an algorithm that uses Bayesian statistical methods to uncover the underlying behavior masked by noisy datasets. This paper outlines the theory behind the SMAUG approach, discusses its implementation, and then uses simulated data and simple experimental systems to show the efficacy of the SMAUG algorithm. Finally, this paper applies the SMAUG method to two model living cellular systems—one bacterial and one mammalian—and reports the dynamics of important membrane proteins to demonstrate the usefulness of SMAUG to a variety of systems.

## Introduction

Super-resolution fluorescence microscopy is a powerful probe of subcellular biology. Single-particle tracking (SPT) measurements have played a central role in measuring the regulation and dynamics of biomolecules inside living cells (1). In both prokaryotes and eukaryotes, SPT measurements have uncovered the positioning and interactions inherent to many different systems, from membrane proteins and lipids to transcription factors and DNA replication machinery (1, 2). By imaging individual fluorescently labeled molecules, determining their sub-pixel positions, and then connecting these localizations into trajectories, one can reconstruct the path of a molecule to measure the physical and biological interactions that govern its motion (3). SPT can measure diffusion coefficients, net velocities, and dwell times; here, we focus on the application of determining the apparent diffusion coefficients of molecules from single-particle tracks.

In this paper, we address a key challenge in single-particle tracking (SPT) analysis: our ability to interpret the data provided by high-quality experimental measurements is limited by the analysis framework. In general, the average apparent diffusion coefficient of a protein can be determined from a linear fit to the mean squared displacement (MSD) as a function of time lag (4). However, a single average diffusion coefficient might no longer be sufficient to describe the system if, for instance, situations where a combination of fast-and slow-moving molecules reflects the free diffusion and the bound state of a molecule, respectively (Fig. 1A). In these cases, the framework can be extended by fitting the MSD vs. time lag to a model for two (or possibly more) diffusive populations (5, 6). Still, curve fitting-based approaches require a predetermined model for the fit, or else a best model can be chosen from a collection given some quality metric such as a penalty function (7) or inspection of residuals (8). Even with a rigorous evaluation of the goodness of fit, these approaches maintain the possibility of user bias in the choice of the pre-approved models and the goodness of fit criteria.

**Figure 1.**
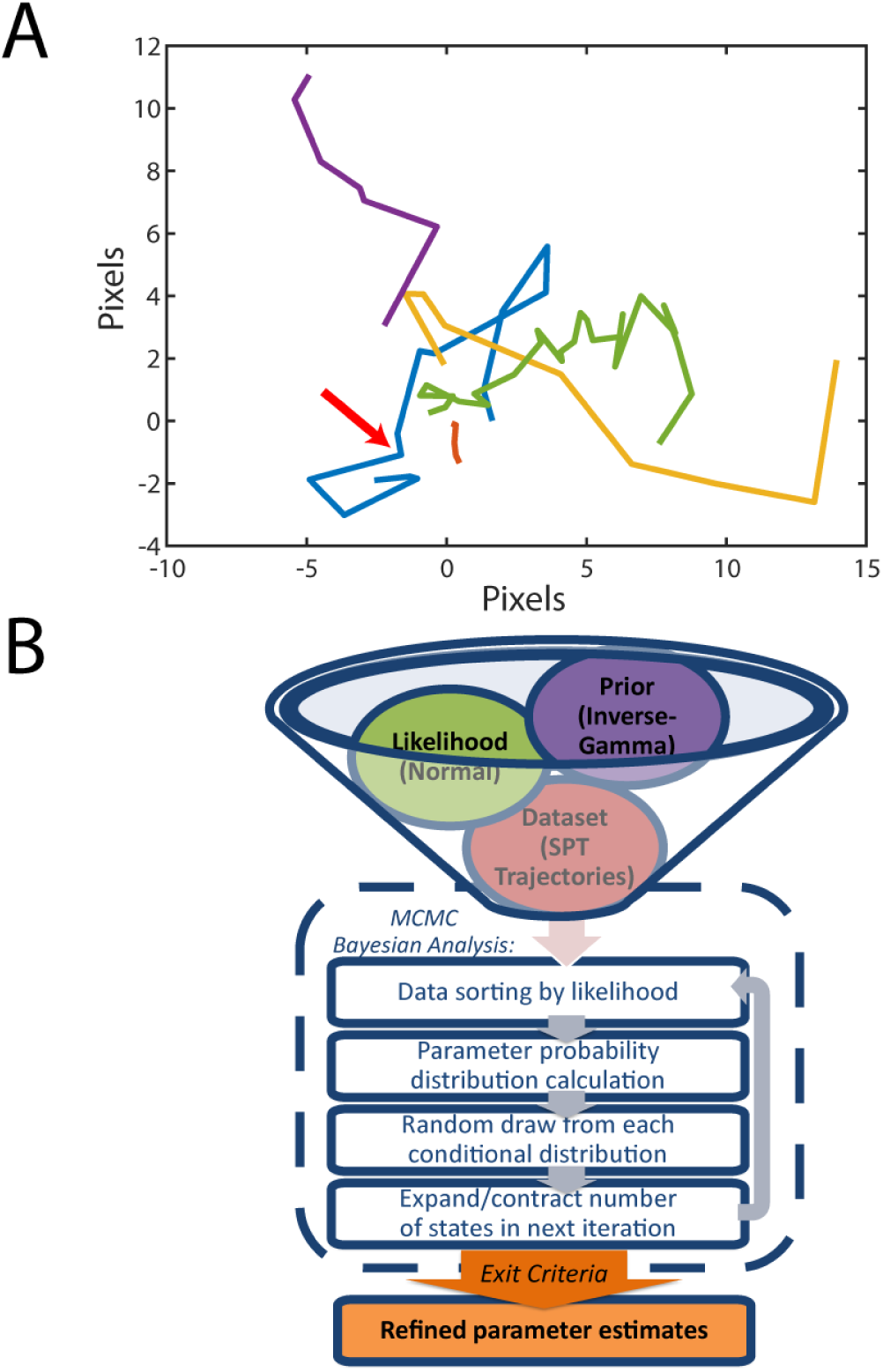
The Single-Molecule Analysis by Unsupervised Gibbs sampling (SMAUG) algorithm. A) A collection of five single-particle trajectories (SPTs) in an environment where single molecules can diffuse at different rates and transition from one state to another. The yellow trajectory has a large average diffusion coefficient, whereas the green trajectory has a small average diffusion coefficient. The red arrow along the blue trajectory marks a transition between states that is difficult to identify by eye. B) Graphical representation of the SMAUG algorithm, which combines the likelihood, prior, and dataset (top) into a Bayesian framework Markov Chain Monte Carlo (MCMC) algorithm that iterates through four steps to refine the parameter estimates (dashed line) until some exit criteria are satisfied.

Here, we address these limitations in the state of the art, and we provide a supervisory-free, hands-off method for measuring heterogeneous single-molecule dynamics by applying nonparametric Bayesian estimation to SPT experiments, with a <u>S</u>ingle-<u>M</u>olecule <u>A</u>nalysis by <u>U</u>nsupervised <u>G</u>ibbs sampling (SMAUG) approach. Bayesian statistical approaches provide a flexible and robust framework for estimating parameter values from experimental data and they offer an alternative to traditional, curve fitting-based techniques. In contrast to these more familiar approaches, which fit data to a function with adjustable parameters, the Bayesian framework does not rely on a pre-determined model. Instead, Bayesian algorithms estimate the most probable parameters by investigating regions in parameter space where the posterior function is very high in order to form a type of topological map of the parameter space. Bayesian approaches have been extensively reviewed, for instance in refs (9–11). Recently, Bayesian analysis techniques have gained popularity in single-molecule biophysics due to their robustness and flexibility. Several recent applications include: analyzing Förster resonance energy transfer (FRET) traces and stepwise photobleaching curves (12), increasing the ability to find and track molecules within single-molecule imaging movies (13), attaining more information from MSD curves of tracked molecules (14), and mapping the local diffusion coefficients within a cell based on the single-molecule trajectories in each small constructed domain (15).

In this paper, we introduce SMAUG, an algorithm that uses Gibbs sampling to implement a nonparametric Bayesian approach to estimate the most probable information about a heterogeneous collection of mobile molecules. In SPT experiments where multiple biochemical functions give rise to multiple observable mobility states, the SMAUG approach allows us to accurately and precisely determine the underlying parameters of the system. In such a system, the biophysical behavior is described by a set of mobility states, each with an average apparent diffusion coefficient and weight fraction, as well as by the likelihood of transitions between mobility states. Here, we use SMAUG to extract these parameters from a collection of single-molecule trajectories free of any supervisory bias such as *a priori* model selection or parameter constraints. Importantly, we uncover novel information that could not be attained with many previous approaches: the probability that a molecule in one mobility state will transition to a different state. The full list of the parameters achieved by SMAUG and a schematic of the algorithm are presented in Table 1 and Fig. 1B, respectively. First, we present the theory of Bayesian inference and its application to SPT. We then validate the SMAUG algorithm on simulated SPT datasets. Finally, we apply SMAUG to SPT experiments *in vitro*, in bacterial cells, and in eukaryotic systems. Overall, SMAUG provides a concrete mathematical framework that can interpret SPT datasets to provide novel biological insight.

**Table 1.** Set of parameters estimated by the SMAUG algorithm to describe a collection of single-molecule trajectories experiencing *K* mobility states.

| Parameter | Symbol | Description |
| --- | --- | --- |
| Number of mobility states | $K$ | Number of distinct mobility states present in the dataset |
| Diffusion Coefficient | $\mathbf{D}$ | $1 \times K$ vector of diffusion values in $\mu\text{m}^2/\text{s}$ for each state |
| Localization Noise | $\epsilon^2$ | $1 \times K$ vector of localization noise for each state |
| Weight Fraction | $\boldsymbol{\pi}$ | $1 \times K$ vector describing the fraction of the data in each state |
| Transition Matrix | $\mathbf{T}$ | $K \times K$ matrix where $K_{ij}$ describes the probability of transitioning from state $i$ to state $j$ |
| Theta | $\boldsymbol{\theta}$ | A vector of vectors that contain all the above information |

## Materials and Methods

### Data Analysis

The analysis algorithms used are described in detail in the Theory section. All code and some test datasets are available as Supporting Material.

### Simulated SPT Trajectories

Simulations of SPT experiments were constructed with custom-built MATLAB code (Matlab R2017b, The MathWorks). Each track was constructed with a random track length drawn from an exponential distribution with mean 10 localizations. Each step along the track could belong to one of several mobility states with corresponding diffusion coefficient, *Di*. Mobility state labels were assigned for each localization by a random draw from the Transition Matrix. Steps along the trajectory were then constructed using a zero-mean Gaussian distribution with variance equal to 2*Di*Δ*t*, where Δ*t* is the frame imaging time. Camera noise and motion blur were applied as described in (16). The “realistic” range imaging parameters were based on reference (8).

### *In vitro* experiments

Fluoresbrite^®^ microspheres with diameters of 100, 200, and 350 nm (Cat # 21636, Polysciences Inc.) in water were diluted 1:1 v/v with glycerol, and 5 µL of the 50% glycerol mixture was placed between two glass coverslips and imaged with a frame exposure time of 40 ms. Imaging was done in an Olympus IX71 inverted epifluorescence microscope with a 60× 1.20 NA water-immersion objective. Samples were excited by a 488 nm laser (Coherent Sapphire 488-50) with power density 140 W/cm^2^. The fluorescence emission was filtered with appropriate filters and imaged on a 512 × 512 pixel Photometrics Evolve electron-multiplying charge-coupled device (EMCCD) camera. Recorded single-molecule positions were detected and localized using home-built code as previously described (5), and connected into trajectories using the Hungarian algorithm (17).

### *Vibrio cholerae* experiments

*V. cholerae* cells containing a chromosomal fusion of the photoactivatable red fluorescent protein, PAmCherry, to TcpP, a membrane-localized transcriptional regulator (TcpP-PAmCherry) as the sole source of TcpP. TcpP-PAmCherry is expressed at the native *tcpP* locus (strain LD51) and cells were grown under conditions known to stimulate TcpP-mediated expression of virulence genes (18) (LB rich media at pH 6.5 and 30 °C). Once cells reached mid log-phase they were diluted into M9 minimal media, and then imaged at room temperature on agarose pads using a 406-nm laser (Coherent Cube 405-100; 102 W/cm^2^) for photo-activation and a 561-nm laser (Coherent-Sapphire 561-50; 163 W/cm^2^) for imaging. Continual images were collected with a 40-ms exposure time per frame in an Olympus IX71 inverted epifluorescence microscope with a 100× 1.40 NA oil-immersion objective. The fluorescence emission was filtered with appropriate filters and imaged on a 512 × 512 pixel Photometrics Evolve EMCCD camera. Recorded single-molecule positions were detected and localized as previously described using home-built code (5), and connected into trajectories using the Hungarian algorithm (17).

### B-cell receptor (BCR) experiments

The BCR dynamics were measured in CH27 mouse lymphoma B cells (RRID: CVCL_7178) as described in (19). Briefly, cells were transiently expressing full-length versions of Lyn kinase or LAT2 (linker for activation of T cells 2)/LAB (linker for activation of B cells) conjugated to mEos3.2. Endogenous, plasma membrane-localized BCR was labeled for 10 min at room temperature with 5 mg/mL goat anti-mouse IgM (Jackson ImmunoResearch; RRID: AB_2338477) f(Ab)1 fragments conjugated to both silicon rhodamine (SiR) dye (Spirochrome, Switzerland) and biotin. Cells were imaged in a live-cell buffer compatible with BCR signaling both before and after the addition of 1µg/ml streptavidin, which clusters and activates receptors. Imaging was performed on an Olympus IX81-XDC inverted microscope with a cellTIRF module, a 100× UAPO TIRF objective (NA = 1.49), and active Z-drift correction (ZDC).

Excitation of the SiR dye was accomplished using a 647 nm solid-state laser (OBIS, 100 mW, Coherent, Santa Clara, CA). Photoactivation of mEos3.2 was accomplished with a 405 nm diode laser (CUBE 405-50FP, Coherent) with excitation using a 561 nm solid-state laser (Sapphire 561 LP, Coherent). All images were taken on an iXon-897 EMCCD camera (Andor, CT) at approximately 45 frames/s with an exposure time of 20 ms. Recorded single-molecule positions were detected, localized, and connected into trajectories as described in (20). Data acquired for Lyn was reported previously (19) and reanalyzed for this work.

## Theory

### Bayesian Statistics

Using SMAUG, we interpret collections of single-particle tracks: a set of single-molecule positions that are connected in time as the molecule moves along some path. The data set, ***y***, is therefore a distribution of step sizes that remain connected by their trajectories, and this data set is the consequence of the physical parameters that govern the single-molecule motion (Table 1):

i. The number of distinct mobility states (modes of motion), ***K***;
ii. The diffusion coefficient, ***D***_*j*_, of each of mobility state;
iii. The inherent localization noise, 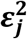, of each mobility state;
iv. The fraction of molecules, ***π***_*j*_, in each of mobility state; and
v. The transition matrix, **T**, where *T*_ij_ is the probability that a step in mobility state *i* precedes a step in mobility state *j*.

Here, we consider these physical parameters as a vector of parameters, ***θ*** = {***D, ε***^2^, ***π***, ***T***}.

Whereas traditional fitting algorithms assume a model function, *f*, and fit the function to the set of parameters, Bayesian estimation calculates the posterior distribution, *p*(*θ*|*y*), which is the probability that a vector of parameter values, ***θ***, gives rise to some data, ***y***. In general, this estimation is accomplished by calculating *p*(*θ*|*y*) according to Bayes’ Rule:

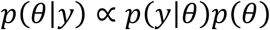

Bayes’ rule relates the posterior distribution, *p*(*θ*|*y*), to the likelihood, *p*(*y*|*θ*), which describes the probability that some parameter vector gives rise to our data, and *p*(*θ*), the prior on our parameters, which encodes our knowledge about the system before any data is collected. In the most general and simple of cases, the posterior is calculated by evaluating the data set, ***y***, at all possible parameter values and then looking for the point at which the calculation is maximized. However, for more complex cases where the posterior can be a mixture of several states and/or where several different posteriors distributions must be calculated, straightforward calculation of all parameter space is impractical if not impossible. In such cases, the main goal of a Bayesian algorithm remains the same: to calculate the posterior in order to find regions of high probability that in turn describe probable values that explain the dataset. However, in these cases of heterogeneous data, the calculation requires methods that are more advanced.

Accordingly, SMAUG enables Gibbs analysis for the applications in this paper by embedding a Gibbs sampling scheme (21, 22) within a Markov Chain Monte Carlo (MCMC) framework (23). SMAUG implements this Markov scheme iteratively in two broad steps (Fig. 1B). In the first step, SMAUG calculates the posterior distribution of the parameters ***θ*** using Gibbs sampling. The Gibbs sampling method iteratively updates each parameter’s posterior individually while holding all other parameters constant to reduce an otherwise impossibly complex posterior calculation into manageable and calculable chunks. In the second step, new parameter values are sampled (i.e., values of ***θ*** are pulled from these calculated posterior distributions) and saved for the first step of the next iteration.

Here, the posterior is a mixture model (more than one mobility state can be present in the dataset), so a data-selection step precedes the sampling step. In this data-selection step, each data point in ***y*** is assigned to a particular mobility state of the mixture, and only the subset of data belonging to that state is used in the posterior calculations that describe that state. Below, we describe our Gibbs sampling process for a known number, *K*, of mixture states; afterward, we describe the process for expanding to an infinite number of states, which is necessary for the sampler to learn the correct number of states present in a dataset.

We can now discuss how SMAUG actually achieves the steps of the Markov Chain process. For a model of given complexity, *K*, we compute the <u>conditional</u> posterior distribution: the posterior distribution for one single parameter while all other parameters remain constant. In this mixture of *K* states, first, we need to assign data to the various states. To identify which data point comes from which of the *K* mixture states, we introduce a latent variable, *l*_*i*_, which labels each of the data points with the number of the state from which it was likely drawn. *L*_*j*_ is the set of all data points with *l*_*i*_ = *j*. For each *l*_*i*_, we calculate the likelihood function, *p*(*y*_*i*_|*θ*_*j*_), (explained below) for each data point belonging to each of the *K* states individually and then we draw the assignment using the categorical distribution with weights equal to the likelihood calculated for each state:

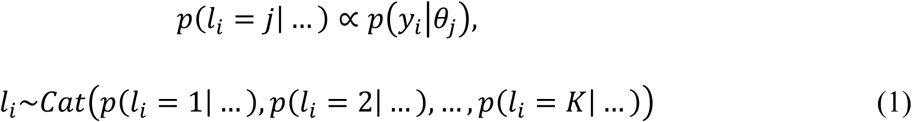

The very first assignment can be random, as information about the likelihood has not yet been calculated. In every iteration, having sorted the data into subsets *L*_*j*_ that are relevant to each state, SMAUG then proceeds to the Gibbs sampler.

Two of the parameters we wish to find for our dataset are the diffusion coefficient values, ***D_j_***, and the localization noise, 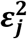. In a model derived by Berglund (16, 24), we relate our dataset of step sizes from SPT experiments to these quantities of interest. By this model, the measured steps sizes, Δ*x*, are zero-mean Gaussian variables whose covariances are related to ***D_j_*** and 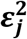 by:

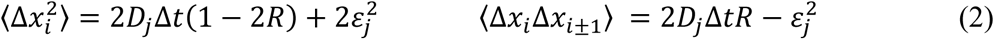

where ∆*t* is the exposure time of the frame and *R* is the motion blur coefficient, which is set to 1/6 as our acquisition and exposure times are equal (16). While SMAUG could be adapted to include other physical manifestations like confinement, we focus here on apparent free diffusion and therefore we model the trajectories as from the result of a zero-mean Gaussian process as stated above. Specifically, the <u>likelihood function</u>, *p*(*y*|*θ*), is a Gaussian denoted *N*(*μ*, *σ*^2^) with unknown mean, *µ*, and unknown variance, *σ*^2^. For most purposes, these step size distributions should be zero-mean (*µ* = 0), but we retain the unknown mean parameter to be as general as possible. Since we have specified that there are *K* distinct states in this dataset, we expand the likelihood to a Gaussian Mixture Model (25) that includes *K* such Gaussian distributions, each scaled by the amount of data in that state, expressed as the fraction of the whole, ***π_i_*** (this weight parameter is discussed more below):

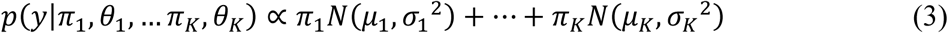

For the <u>prior distribution</u>, SMAUG takes the conjugate prior to our likelihood: the Inverse-Gamma function, *IG*(*a, b*). A conjugate prior is a prior distribution that when multiplied to the likelihood returns a posterior that is of the same mathematical family as the likelihood, simplifying the computation. By constructing the likelihood and the prior this way, SMAUG arrives at the full <u>conditional posterior function</u> for diffusive motion: the Normal-Inverse-Gamma function, *NIG*(*μ_K_*, *σ*_*K*_^2^, *a*, *b*):

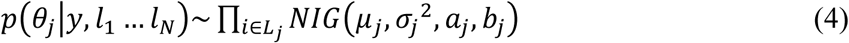

The conditional posterior that describes the weight fractions, ***π***_***j***_, for each state is a standard Dirichlet distribution, *DIR.* The Dirichlet is a generalization of the Beta distribution and always sums to 1. Using the set of *L*_*j*_ as inputs:

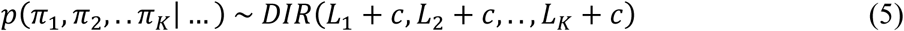

where the constant vector [*c,…,c*] is used with the conjugate Dirichlet distribution (the Dirichlet distribution is conjugate to the categorical distribution and results in a Dirichlet posterior) to describe the prior weight. Transitions between states are also sampled from the Dirichlet distribution. Using the assignment values of *l*_*i*_ from before, SMAUG creates a transition matrix that measures when subsequent steps within a trajectory change their assignment. *N*_a,b_ counts the number of transitions from state *a* to state *b*. Each of the rows of the transition matrix, ***T***, are sampled as:

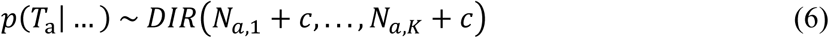

where the vector [*c,…,c*] acts as the prior weight vector.

Thus, for any mixture of *K* states, the Gibbs scheme outlined above efficiently samples from the conditional posteriors of all the model parameters. In the second step of the Markov Chain, these newly defined distributions are sampled to get the parameter values that will be used in the calculations of the next iteration. However, we rarely know *a priori* how many distinct states to include in an analysis; in fact learning this number can be one of the principle goals of an SPT experiment (5, 8, 26).

We could set up some large upper bound for *K*, but then much of our computational power would be directed towards calculating states with zero occupancy, leading to a computationally inefficient process. Instead, in the next section, we outline a Dirichlet process mixture model (DPMM) method that allows the number of states to expand or contract organically in response to the data based on a nonparametric Bayes approach.

### Nonparametric Bayes

Nonparametric Bayesian techniques rely on random probability measures to extend a finite-component mixture like the one described above into an infinite-component mixture model needed for completely hands-off estimator (free from supervisory bias) (27). SMAUG uses the Dirichlet process, DP(*α*, *P*_0_), one such random probability measure which is generally described as a “distribution of distributions”. Specifically, DP(*α*, *P*_0_) is a distribution with base probability distribution, *P*_0_ (such as a normal Gaussian or a beta distribution), and concentration parameter, *α* (which controls the variance around *P*_0_). The DP can be seen as the infinite dimensional generalization of the standard Dirichlet distribution and, as with the standard Dirichlet, the “weights” drawn must sum to 1, which helps induce a clustering onto the infinite collection of possible states present in the nonparametric realization of the sampler. A helpful visualization for understanding what a draw from a Dirichlet process looks like is the stick-breaking construction (28), which represents the total probability available to the system as a stick of unit length. First, a random sample, *θ*_1_~*P*_*0*_, is drawn from the base probability measure *P*_0_ (*θ*_1_ can be a single value or a vector), and random weight, *V*_1_ ~ Beta(1, *α*), is pulled from the distribution. We give a probability weight of *π*_1_ = *V*_1_ to point mass *θ*_1_. We then break our unit stick at *V*_1_ and there now remains an amount of stick, (1 – *V*_1_), to be allocated to the many other draws. We then break an amount *V*_2_ ~ Beta(1, *α*) off the remaining stick and assign probability *π*_1_ = *V*_2_ (1 – *V*_1_) to another point mass of probability *θ*_2_~*P*_0_. As we continue, the stick gets shorter and shorter and so the weight assigned to each new draw from *P*_0_ decreases with a rate that depends on the concentration parameter, *α*. Thus, our random probability measure, *P*, is an arbitrarily large collection of segments of which only several have the vast majority of the probability weight; the rest of the segments have negligible mass. This stick-breaking construction can be summarized as:

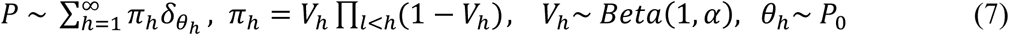

where the *θ*_h_ parameter value vectors are generated independently from *P*_0_, 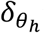 is the point mass where the parameters *θ*_h_ are concentrated, and *π*_h_ is the probability mass associated with that point mass.

SMAUG uses the Slice Sampler method from Walker to reduce the infinite state model that results from the distributions above, in Equation (7), to a model with only finitely many states capable of being calculated at each iteration (29). The Slice Sampler method introduces another latent variable, *u*, which is drawn from the uniform distribution as 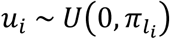, for each data point in the set. Thus, any draw for *u* splits the infinite set of possible states into two categories: a finite set of states for which *π*_j_ > *u* and an infinite set for which *π*_*j*_ < *u* for each data point. By looking for the minimum entry of the set of all *u* and looking at the size of the finitely many states with probability weight greater than that value we know the maximal size of the model we need to include for any iteration. Specifically SMAUG attempts in every iteration to satisfy the inequality:

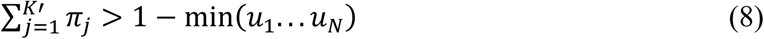

where *K*′ is the number of states present in the model at any time. In this way, only *K*′ states need to be calculated, but over the course of sampling, we integrate over an essentially infinite (or at least arbitrarily large) number of possible states. The value of *K*′ can expand or contract over the course of the analysis with new terms being added when needed and terms whose occupancy is very low (i.e., states of a few data points or less) removed.

Taken together, SMAUG provides an efficient nonparametric Bayesian analysis framework for analyzing SPT data that returns accurate and precise estimates of the number of mobility states within a dataset, their diffusion coefficients, weights, and transitions in a hands-free manner. During each iteration of the sampler, SMAUG follows a simple stepwise process as outlined above (Fig. 1B):

1. **First iteration only**: choose an initial number of states. This number can be selected to be several times bigger than the expected number for the experiment. Assign each of the data points to a state by some method.
2. Assign a vector of parameter values to each state, for instance by random draws from a base distribution, *P*_0_, or by calculating the simple statistics (mean and variance) from the previous step’s assignment.
3. **Second iteration onward**: Assign each data point a latent variable, *u*, and use these values of *u* to determine *K*′, the number of states present for this iteration.

a. If states need to be added, assign each of them a weight by breaking the stick and pulling a parameter vector from *P*_0_.
b. If a state needs to be removed, remove the state with smallest weight and add its weight to the next smallest weight.
c. If states drop out by receiving zero weight fraction, remove the values for that state from the parameter vector.
4. Implement a Gibbs Sampler with the fixed number of states, *K*′, from step 3. Assign labels and update parameters by calculating the conditional posterior distributions described in equations (2–6) above, then sample from these distributions to collect new parameter values for the next iteration.
5. **Exit criteria:** Repeat steps (3) and (4) until some cutoff criterion has been achieved, either based on performing some number of total iterations or attaining some convergence metric, then construct parameter estimates from the back half of saved iterations.

The efficient SMAUG algorithm we have built provides a flexible method for determining all the relevant parameters for an arbitrary SPT trajectory dataset without supervisory bias. For instance, the amount of data generated in step (4) can be controlled by not saving the parameters of interest every iteration (by default, SMAUG saves every tenth iteration to minimize any possible autocorrelation between iterations).

We demonstrate below that SMAUG accurately and precisely estimates for SPT experiments the number of mobility states, the diffusion coefficients, the weight fractions, the noise values, and the frequencies of transitions between states. To demonstrate the value and feasibility of this nonparametric Bayesian algorithm, we validate our method first by using simulated diffusion trajectories with realistic parameter values and an *in vitro* experimental system, and then we apply SMAUG to subcellular tracking in bacterial cells and in eukaryotic cells.

## Results and Discussion

Single-particle tracking (SPT) analyzes the trajectories collected from sequential molecule locations. These trajectories describe the motion of the molecule. However, biological complexity creates heterogeneities even along single tracks, and the SPT trajectories must be carefully considered to yield quantitative information. Previously, we analyzed SPT data by curve-fitting the squared step-size distribution to a cumulative probability distribution (CPD) equation for a model of heterogeneous diffusion (5, 8), and we expanded the number of terms in the model until fits to the data were deemed sufficient by inspection of the residuals subject to a penalty function to limit over-fitting (since residuals will always decrease when a model is expanded).

However, such fitting-based techniques suffer from weak parameter identifiability: generally, several equally acceptable fits can be obtained. To select between these similar outcomes, parameter values can be more tightly constrained based on justifiable knowledge of the system beforehand, which is very rarely available. Furthermore, this approach leaves much of the information from the data set unused, as it decouples the individual steps from their trajectories: CPD methods usually assume that each squared step size is independent, and they ignore the longer trajectories that can give each step context and inform about transitions between states. Here, we improve this data analysis by assuming that each step is a time point in a Markov-like process, in which each subsequent step depends only on the current time point, and we use a nonparametric Bayesian process to discover the underlying parameters using the whole, unmodified dataset.

### Validation of SMAUG with simulated data

We validated the SMAUG algorithm with a simulated dataset (Table S1) containing 13,636 steps (1090 trajectories) drawn from a diffusive mixture with four distinct mobility states, ***i*** = {1,2,3,4}. The diffusion coefficients for the terms were *D* = {0.005, 0.03,0.09,0.20} μm^2^/s, and the localization errors for each localization, 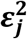, were pulled from a distribution with a mean of 10 nm and a variance of 10 nm. The weight fractions of each term were: {*π*_1_, *π*_2_, *π*_3_, *π*_4_} = {0.196, 0.301, 0.291,0.212}. The transition matrix was:

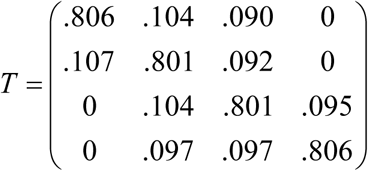

Where *T*_*ij*_ is the probability of a step in mobility state *i* preceding a step in mobility state *j*. In these realistic simulations, the track lengths and transitions are random events: each track length is pulled from an exponential distribution with a mean of 10 localizations, and *t*_*i→j*_, the likelihood of transitioning from state *i* to state *j*, is governed by a flat distribution. The seed values for the simulations, which gave rise to the simulated values, are given in Table S1.

As opposed to methods that fit data to a specific model with a selected number of mobility states, *K*, one strength of the SMAUG algorithm lies in its ability to identify the correct number of mobility states. The algorithm was initialized with a large number of components (*K* ≥ 10), but quickly collapsed to the correct number (*K* = 4) (Fig. 2A). In general, the model complexity is increased or decreased until it converges at the correct number, though SMAUG continues to explore state space by adding states on occasion and then removing them (e.g., the bumps up to 5 in Fig. 2A). To construct parameter estimates for this case, we only use posterior draws from saved iterations where *K* is the convergence value (here *K* = 4) in the back half of all saved iterations (iterations 501-1000, red box in Fig. 2B).

**Figure 2.**
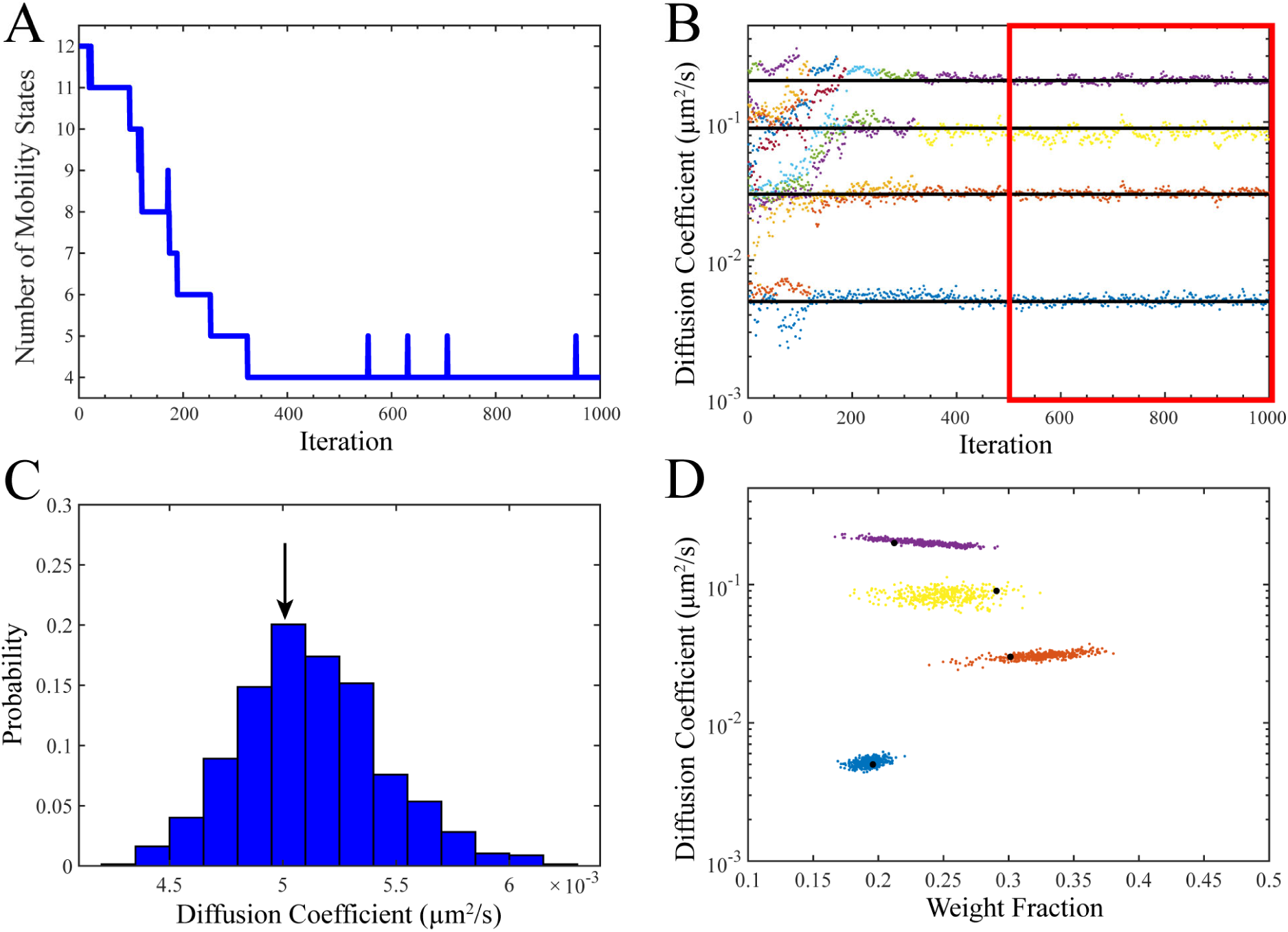
SMAUG Analysis of Simulated Test Data. A) SMAUG analysis of the simulated input data as a Gaussian Mixture Model. The algorithm initializes at a large number of states and quickly converges to the correct value of *K* = 4. However, after convergence additional states are added stochastically as the algorithm explores state space looking for other regions of high probability. B) Estimates of the diffusion coefficients, *D*, for each term (sorted in order of *D*) as the algorithm progresses. Black lines are the true simulation values (Table S1). C) Histogram defining the probability of a given diffusion coefficient for the slowest term in the analysis (term 1 in Table S1). Histograms are constructed using the back half of saved iterations for the blue/slowest term (red box in ‘B’). Black arrow is the true value for the simulation. D) Diffusion Coefficients and weight fraction estimates for each saved iteration in the back half of the analysis run that also meets the *K* = 4 criterion. The analysis shows distinct clusters whose estimates do not overlap. Black dots are the true simulation values. Full histograms for all output values are in SI Fig. S1.

Each parameter is observed throughout the course of the simulation (Fig. 2B, SI Fig. S1) and the terms are sorted by *D* in the final output. We use only the second half of saved iterations (red box in Fig. 2B) to construct estimates. The posterior distributions are plotted for several parameters (Fig. 2C, SI Fig. S1). Because these data points represent draws from the converged posterior distributions for the parameters, we use these histograms to calculate statistics about our estimates or construct confidence intervals. The mean values in all cases are close to the true values (black arrows in Fig. 2C and SI Fig. S1). At each step in the analysis, the best estimates for all parameters are generated, and each pair {*Di*, *π*_*i*_} for every saved iteration is plotted as a point in Fig. 2D. True values for the simulation (Table S1) are indicated by the large black data points in Fig. 2D.

To examine the ability of SMAUG to detect rare occurrences, we simulated a dataset (Table S2) containing 9,445 steps in which the majority of trajectories (95.5 %) belonged to a fast diffusing state (*D*_1_ = 0.15 μm^2^/s) while the rest belonged to a slower state (*D*_2_ = 0.01 μm^2^/s). Furthermore, the transitions between states 1 and 2 were rare (*T*_12_ = *T*_21_ = 0.01). This distribution is relevant for experiments in which the binding events of biomolecules are rare, and analyzing this simulation explores the ability of SMAUG to confidently distinguish rare states from random events within a homogeneous distribution. SMAUG isolates the two distinct populations (SI Fig. S2) and accurately estimates their parameter values (Table S2). SMAUG can easily identify states whose occupancy is only a small fraction of the whole dataset.

### Validation of SMAUG *in vitro*

We further tested the SMAUG method with an *in vitro* experimental system consisting of three different sizes of diffusing fluorescent beads in a 50/50 water/glycerol mixture. The Stokes-Einstein equation predicts a diffusion coefficient of *D* = *KT*/6*πηγ* for a particle of radius, *r*, undergoing Brownian motion in a fluid with viscosity, *η*; this equation predicts theoretical diffusion coefficients of *D* = {0.182, 0.319, 0.637} μm^2^/s for this system. SMAUG analysis of the bead trajectories (Table S3) correctly identified the number of distinct diffusors (*K* = 3) and estimated values of *D* = {0.168, 0.329,0.675} μm^2^/s (SI Fig. S3). The distributions of the estimations of *D* at every saved iteration (SI Fig. S3) show that the theoretical *D* values are within the confidence intervals of the estimations. Furthermore, the transition matrix shows negligible transitions between states (*T*_*ij*(*i≠j*)_ < 0.03). This observation is consistent with our attribution of each state to one bead size as beads cannot change sizes spontaneously and thus no transitions are allowed.

Overall, SMAUG accurately determines the values for parameters of interest (Table S3): the number of distinct mobility states within a dataset; the diffusion coefficient, and the weight fraction, of each state; and the transition probabilities between the states at each iteration for these *in vitro* experiments.

### Application to measuring protein cooperativity in living bacterial *Vibrio cholerae*

We extended SMAUG to live-cell single-molecule tracking to quantify the diffusion coefficients and distributions in biological systems. The pathogenic bacterium *Vibrio cholerae* remains a global health concern, infecting millions each year leading to the diarrheal disease cholera (30). The cholera toxin (CtxAB) and an adherence organelle called the toxin-coregulated pilus (TcpA-F) are key determinants of virulence that are under the regulatory control of ToxT, the expression of which is regulated by the membrane protein TcpP (18). In collaboration with other membrane proteins (TcpH, ToxR, and ToxS), TcpP initiates the *V. cholerae* virulence cascade by binding to the promoter region of the *toxT* gene while remaining in the membrane (Fig. 3A). We are investigating this unusual membrane-localized mechanism of transcription using SPT to measure TcpP dynamics in living cells. Previously, we used fusions to the photoactivatable fluorescent protein PAmCherry to show that TcpP-PAmCherry diffuses heterogeneously in living cells (8).

**Figure 3.**
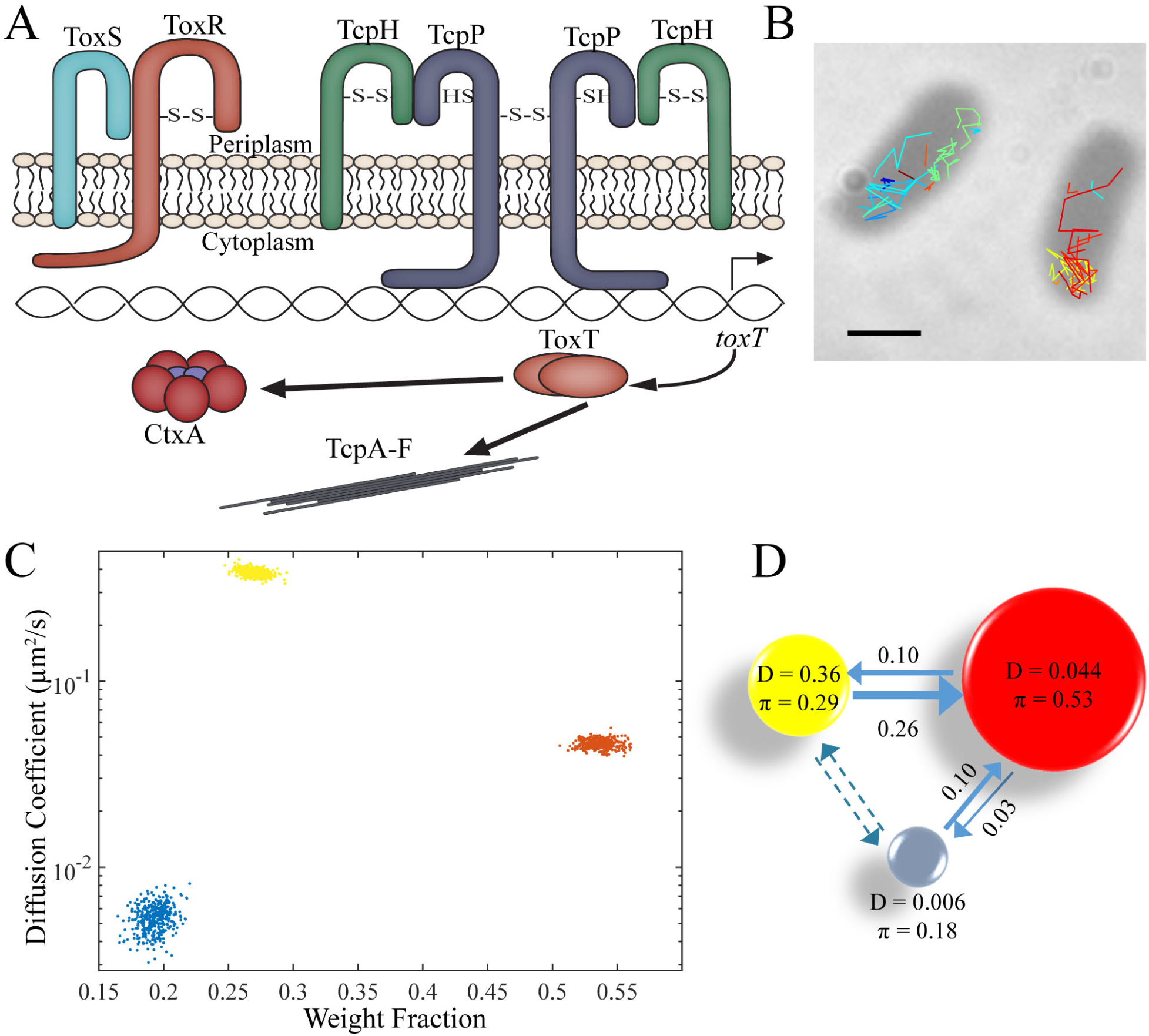
SMAUG analysis for bacterial imaging. A) Schematic of the *V. cholerae* virulence pathway. The membrane-bound protein TcpP binds the DNA directly along with other supporting proteins, leading to the hypothesis that the dynamics of TcpP reflect multiple mobility states. B) Representative image of individual TcpP-PAmCherry molecules trajectories inside live *V. cholerae* cells. Scale bar: 1 µm. C) Diffusion coefficient and weight fraction estimates from the output of the SMAUG analysis. SMAUG identifies three distinct clusters within the dataset. D) Cartoon depiction of the full SMAUG results for this dataset, including transition probabilities. Bubble colors correspond to the term colors in ‘C’ and bubble sizes represent the weight fractions. Arrows between bubbles indicate the mean of the transition matrix elements for transitions between those terms. Dashed lines indicate transition probabilities that are less than 1%.

We tested the SMAUG algorithm on *V. cholerae* cells that encode a chromosomal copy of *tcpP-PAmCherry* that remains under the control of its native promoter (Methods). These TcpP-PAmCherry fusions are functional based on expression levels of downstream protein CtxB (SI Fig. S4) and the fact that the cells (strain LD51) exhibited wild-type growth rates. Furthermore, we observed regular cell morphology under the microscope (Fig. 3B). We grew these cells under virulence-inducing conditions (Methods) and collected 11,403 steps from 2404 trajectories; representative trajectories are shown in Fig. 3B. Analysis of this dataset by SMAUG indicated a most probable interpretation of a *K* = 3-term model with diffusion coefficients of *D*_i_ = {0.006, 0.044,0.368} μm^2^/s and weight fractions of *π*_i_ = {0.18,0.53,0.29} (Fig. 3C – D, Table S4). The combined dataset for TcpP-PAmCherry trajectories results from four days of experiments in 111 cells. We then created 100 independent analysis runs using random sampling with replacement of the entire *V. cholerae* dataset of tracks and found that *K* = 3 was by far the most likely outcome (77 of the runs returned a 3-state model as the most likely (Table S4)).

Figure 3D summarizes the key results for our measurements of TcpP-PAmCherry mobility: we observe three distinct mobility states, which we attribute to different binding states of the protein. Of the three states identified in our experiments, the intermediate state (*D*_2_ = 0.044 µm2/s; red circle in Fig. 3D) is the most highly occupied state (*π*_2_ = 0.53). TcpP exists in the membrane as either a monomer or a dimer (31), and we propose that the fastest diffusive state is free monomeric or dimeric TcpP-PAmCherry (yellow circle in Fig. 3D). We further hypothesize that TcpP association with other proteins in the membrane—most importantly its interaction partners, TcpH and ToxR—leads to its scanning the DNA for its binding target in the *toxT* promoter (8, 18). We propose that the intermediate state is this protein complex DNA-searching state (red circle in Fig. 3D). Finally, to initiate the virulence cascade, this protein complex stops scanning and binds more tightly to the *toxT* promoter region of the DNA, and we propose the slowest term (blue circle in Fig. 3D) is this promoter-bound state. Our model is further supported by the transition matrix (arrows in Fig. 3D and Table S4), which shows negligible transitions from the fastest to the slowest terms and instead outlines a path from the fastest state through the intermediate state to the slowest state, indicating that the TcpP monomers and dimers cannot directly bind the DNA, but rather that TcpP must form a complex with ToxR and/or TcpH before binding DNA and promoting *toxT* transcription. Testing these hypotheses to definitively assign the true nature of these identified mobility states will require further study, and this analysis illustrates the utility of SMAUG for bacterial systems and provides a baseline to which future studies can be compared to more fully understand the mechanistic behavior of the *V. cholerae* virulence mechanism.

### Application to antigen response in eukaryotic B cells

Finally, we applied our analysis method to investigate the dynamics of proteins involved in B-cell receptor (BCR) signaling. Situated in the plasma membrane of B cells, the BCR recognizes and binds antigens, causing BCR clustering and initiating a downstream signaling pathway that results in BCR endocytosis and antigen processing. Following receptor clustering, the BCR is phosphorylated by the Src-family kinase Lyn (32), leading to recruitment and activation of the cytoplasmic kinase Syc which plays multiple roles in propagating the initial immune response. One target of Syc phosphorylation is the transmembrane adaptor protein LAB/LAT2, one of many proteins found within the BCR signalosome (30), a collection of proteins that localize, stabilize, and extend sites of BCR activation. Previously, it was found that membrane domains and lipid organization play a role in BCR activation by clustering BCR receptors upon antigen binding (19).

Using simultaneous two-color super resolution imaging, we analyzed the single-molecule trajectories of BCR and downstream protein Lyn or LAT2 at room temperature before and after stimulation by antigen addition (19) (Fig. 4A – B). We split the trajectories into groupings of 1000 frames; each group contained on average 20,000 – 30,000 steps and occurred over ~22 s, during which time frame we assume the dynamics do not change. In this way, we used SMAUG to analyze the evolution of the dynamics of the system over time. Before stimulation (Fig. 4C, left and Fig. 4D, first bar), the BCR dynamics are best described by three mobility states, with very little weight fraction in the slowest state (red). In other words, most BCR molecules are highly mobile. Immediately after stimulation (Fig. 4D, second bar), SMAUG finds four mobility states: the intermediate term is split into two mobility terms (brown and yellow). This finding may indicate a transition shortly after stimulation. Quickly, SMAUG returns only two mobility states, one of which is not observed pre-stimulation (blue) which we attribute to a new physiological state (SI Fig. S5). The most mobile terms have disappeared from the analysis as the system responds to antigen stimulation. This slower collection of mobility states persists for several minutes until the end of the measurement.

**Figure 4.**
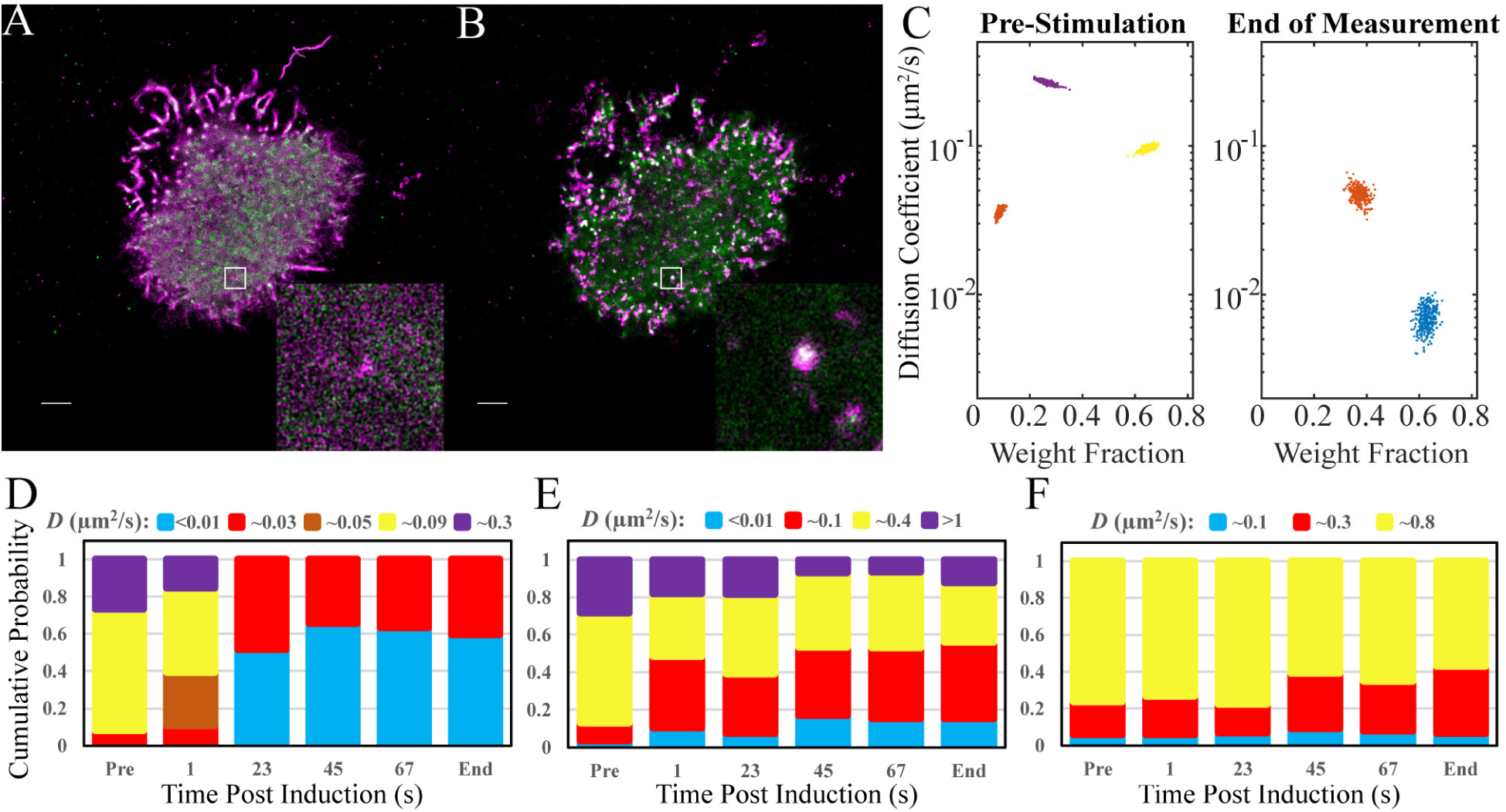
SMAUG analysis for single-molecule motion in a eukaryotic system. A) Super-resolution reconstruction image of BCR-SiR (magenta) and LAT2-mEos3.2 (green) in a representative B cell pre-stimulation. White: overlapping magenta and green signals. Inset is a higher resolution reconstruction of the 1.5 µm × 1.5 µm white boxed region. Scale bar: 2 µm. B) Super-resolution image of the cell in ‘A’ 12.8 min post-stimulation. White: overlapping magenta and green signals. Inset shows same 1.5 µm × 1.5 µm white boxed region as in ‘A’ at a higher resolution. Scale bar: 2 µm. C) Diffusion coefficient and weight fraction estimates for BCR molecules pre-stimulation and at the end of the measurement. Three distinct clusters are found pre-stimulation, but only two at the end of the measurement. D) Bar graphs showing the mean weight fraction of each identified state as a function of time for the BCR dataset. The bars labeled “Pre” and “End” correspond to the data in ‘C’. All other bars are labeled with the time post-stimulation. Identified mobility states are states whose estimates overlap in diffusion coefficient and weight fraction. A new, slower state (blue) emerges 23 s after antigen stimulation. E) The bar graphs for the weight fractions of LAT2 states over time show that the slowest mobility states (blue and red) increase in weight fraction relative to the faster terms (yellow and purple) suggesting the assembly of the BCR signalosome. F) The bar graphs for the weight fractions of Lyn states over time show that there is no change upon antigen stimulation and a slight overall decrease in mobility of the system starting at ~45 seconds. The full cluster analysis is in SI Fig. S5.

Simultaneously, we monitored the dynamics of Lyn or LAT2, and we matched the dynamics of the downstream protein with the response from the BCR itself. Analysis of LAT2 indicates four mobility states whose dynamics change greatly after BCR stimulation (Fig. 4E). Like BCR, the LAT2 dynamics slow over time post-stimulation: the slower LAT2 mobility state’s population fraction increases and the faster LAT2 mobility state’s population fraction decreases after stimulation. In contrast, for Lyn, a tyrosine kinase and the first protein in the downstream cascade, analysis with SMAUG consistently returns a three-term model with similar weight fractions and diffusion coefficients before and after BCR stimulation (Fig. 4F), with a slight change in the weight of the middle term occurring at ~45 seconds and persisting through the end of the measurement. Consistent with this mobility analysis, we find that LAT2 colocalizes much more strongly with cross-linked BCR than does Lyn. A second analysis on different cells returns very similar results to those described above (SI Fig. S6). More studies are needed to assign biochemical and biophysical roles to the states uncovered by SMAUG, but this experiment proves the efficacy and utility of SMAUG analysis for both eukaryotic systems as well as for time series data.

## Conclusions

Single-molecule experimental techniques have greatly enhanced the field of biophysics and our understanding of many biological problems. However, as SPT experiments are extended to include more complex systems, the need for a mathematically rigorous analysis method has increased. The SMAUG method we developed in this paper allows completely hands-free analysis of single molecule tracking data by using a nonparametric Bayesian approach to fully characterize the posterior distributions of many of the relevant parameters and to quantify the corresponding parameter uncertainties. Such a method allows more concrete and objective conclusions to be drawn from SPT experiments as it bypasses the issues of supervisory bias and model selection that can alter the data processing and the conclusions drawn. In this paper, we have validated the method using realistic simulations and *in vitro* systems and then displayed this method’s ability to uncover hidden information in a biological system by investigating an open question in biology in both prokaryotes and eukaryotes. The SMAUG analysis method attempts to provide researchers with a powerful and easy to use tool for analyzing SPT experiments and drawing concrete conclusions to provide insight into biomolecule dynamics in many relevant systems.

## Supporting information

Supplemental - SMAUG Code

## ASSOCIATED CONTENT

### Supporting Information

Supporting Tables S1 – S4 and Supporting Figures S1 – S6

### Code Availability

Open-source Matlab code for implementing SMAUG (GNU General Public License) and some test datasets are provided (Supplementary Information). Further development and expansion of the code post-publication will be hosted at https://github.com/BiteenMatlab/SMAUG.

## Author Contributions

J.D.K. and J.S.B. designed the research. J.D.K. implemented the SMAUG algorithm, wrote the code, performed the simulations, and analyzed the data. J.D.K. and E.D.D. performed the *in vitro* and bacterial imaging experiments. S.A.S. performed the B cell imaging experiments. L.M.D. constructed the bacterial strains and performed the biochemical assays. All authors discussed the results. The manuscript was written and edited by all authors. All authors read and approved the manuscript.

## ACKNOWLEDGEMENTS

This work was supported by a National Institutes of Health award (Grant 1-R21-GM128022-01) to JSB and R21-AI099497 to VJD and JSB and R01GM110052 to S.V. Thanks to Stephen Lee and Jason Karslake for helpful discussions.

---

**Table S1.** Seed values, true values, and SMAUG results for the four-term simulation described in Figs. 2 and S1. Total number of steps included is 13,636.

| Parameter | Seed Values | True Values | SMAUG Results |
| --- | --- | --- | --- |
| Number of Mobility States | 4 | 4 | 4 |
| Diffusion Coefficients ($\mu\text{m}^2/\text{s}$) | {0.005, 0.03, 0.09, 0.2} | {0.005, 0.03, 0.09, 0.2} | {0.0051, 0.0305, 0.836, 0.201} |
| Standard Deviation | N/A | N/A | {0.0003, 0.0016, 0.0082, 0.0093} |
| Localization Noise (nm) | $10 \pm 5$ | $10 \pm 5$ | {5.7, 8.8, 9.6, 13.7} |
| Weight Fractions | {0.25, 0.25, 0.25, 0.25} | {0.196, 0.301, 0.291, 0.212} | {0.192, 0.322, 0.251, 0.235} |
| Standard Deviation | N/A | N/A | {0.0076, 0.0223, 0.0234, 0.0235} |
| Transition Matrix | $\begin{pmatrix} 0.8 & 0.1 & 0.1 & 0 \\ 0.1 & 0.8 & 0.1 & 0 \\ 0 & 0.1 & 0.8 & 0.1 \\ 0 & 0.1 & 0.1 & 0.8 \end{pmatrix}$ | $\begin{pmatrix} .806 & .104 & .090 & 0 \\ .107 & .801 & .092 & 0 \\ 0 & .104 & .801 & .095 \\ 0 & .097 & .097 & .806 \end{pmatrix}$ | $\begin{pmatrix} .812 & .132 & .033 & .023 \\ .069 & .776 & .124 & .031 \\ .023 & .166 & .676 & .134 \\ .022 & .047 & .145 & .785 \end{pmatrix}$ |

**Table S2.** Seed values, true values, and SMAUG results for the rare-states simulation in Fig. S2. Total number of steps included is 9,445.

| Parameter | Seed Values | True Values | SMAUG Results |
| --- | --- | --- | --- |
| Number of Mobility States | 2 | 2 | 2 |
| Diffusion Coefficients ($\mu\text{m}^2/\text{s}$) | {0.01, 0.15} | {0.01, 0.15} | {0.011, 0.146} |
| Standard Deviation | N/A | N/A | {0.0011, 0.0019} |
| Localization Noise (nm) | $10 \pm 10$ | $10 \pm 10$ | {12.1, 15.7} |
| Weight Fractions | {0.05, 0.95} | {0.045, 0.955} | {0.048, 0.952} |
| Standard Deviation | N/A | N/A | {.0033, .0035} |
| Transition Matrix | $\begin{pmatrix} 0.99 & 0.01 \\ 0.01 & 0.99 \end{pmatrix}$ | $\begin{pmatrix} 0.987 & 0.013 \\ 0.009 & 0.991 \end{pmatrix}$ | $\begin{pmatrix} 0.969 & 0.031 \\ 0.001 & 0.999 \end{pmatrix}$ |

**Table S3.** Theoretical values and SMAUG results for the diffusing beads experiments in Fig. S3. The theoretical diffusion coefficient is calculated from the Stokes-Einstein Equation. The theoretical weight fraction is based on taking the fraction of number of steps that came from each bead in the total combined data set. The theoretical transition matrix includes no transitions as the beads cannot spontaneously change sizes. Total number of steps included is 31,949.

| Parameter | Theoretical Values | SMAUG Results |
| --- | --- | --- |
| Number of Mobility States | 3 | 3 |
| Diffusion Coefficients ($\mu\text{m}^2/\text{s}$) | {0.182, 0.319, 0.637} | {0.168, 0.329, 0.675} |
| Standard Deviation | N/A | {0.0027, 0.0083, 0.0166} |
| Weight Fractions | {0.411, 0.350, 0.239} | {0.423, 0.358, 0.219} |
| Standard Deviation | N/A | {0.0110, 0.0119, 0.0106} |
| Transition Matrix | $\begin{pmatrix} 1 & 0 & 0 \\ 0 & 1 & 0 \\ 0 & 0 & 1 \end{pmatrix}$ | $\begin{pmatrix} 0.977 & 0.018 & 0.005 \\ 0.021 & 0.946 & 0.036 \\ 0.008 & 0.059 & 0.932 \end{pmatrix}$ |

**Table S4.** Summary of measurements for TcpP-PAmCherry mobility in *V. cholerae* (Fig. 3). Total number of steps in the original dataset is 11,403.

| Parameter | SMAUG Results<br>(All data) |  | Parameter | Bootstrap Results* |
| --- | --- | --- | --- | --- |
| Number of mobility states | 3 | | Number of runs with $K = 3$ | 77% |
| Diffusion Coefficients ($\mu\text{m}^2/\text{s}$) | {0.0054, 0.046, 0.383} | | Mean Diffusion Coefficients<sup>§</sup> ($\mu\text{m}^2/\text{s}$) | {0.0053, 0.043, 0.355} |
| Standard Deviation | {0.0009, 0.0025, 0.0168} |  | Standard Deviation | {0.0011, 0.0043, 0.0181} |
| Weight Fractions | {0.193, 0.537, 0.270} |  | Mean Weight Fractions<sup>§</sup> | {0.171, 0.540, 0.289} |
| Standard Deviation | {0.0095, 0.0094, 0.0079} |  | Standard Deviation | {0.0195, 0.0164, 0.0133} |
| Transition Matrix | $\begin{pmatrix} .839 & .122 & .039 \\ .046 & .810 & .145 \\ .025 & .291 & .684 \end{pmatrix}$ | | Mean Transition Matrix<sup>§</sup> | $\begin{pmatrix} .872 & .108 & .020 \\ .042 & .799 & .159 \\ .016 & .329 & .655 \end{pmatrix}$ |

\*Bootstrapping was performed over 100 rounds. 6% of analysis rounds yielded a 2-state model, 77% gave 3 states, 14% gave 4 states, and 3% gave 5 states.
§ Mean and standard deviation values are constructed by using only the runs where $K = 3$ . A new vector was created from the mean outputs of each of the 77 individual runs, and the mean and standard deviation of that vector are reported here.

**Figure S1.**
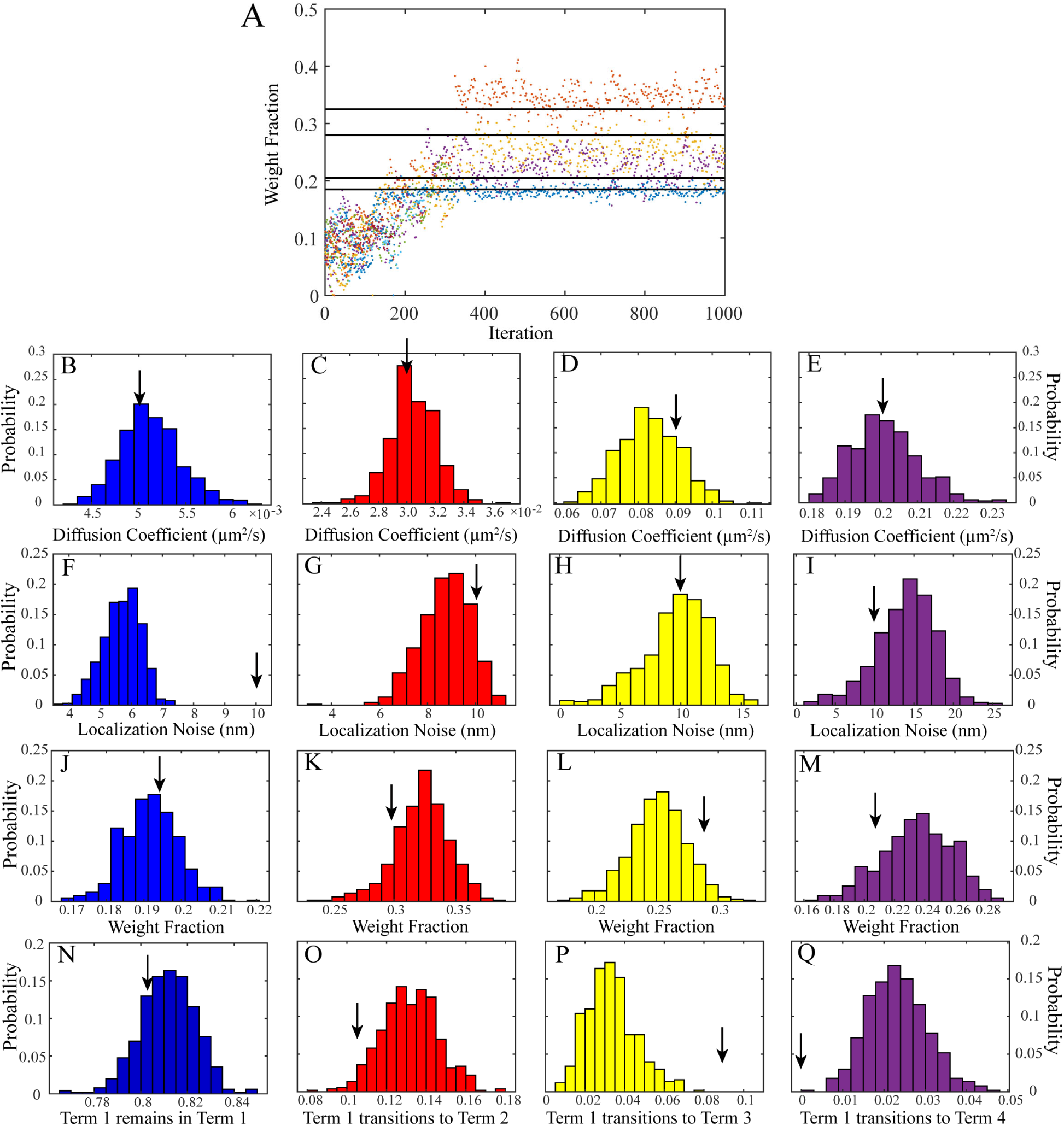
Full SMAUG analysis for simulated data. A) Estimates of the weight fraction for each term (sorted in order of increasing diffusion coefficient) as the algorithm progresses. Black lines are the simulation true values (Table S1). B – E) Histograms of the diffusion coefficient estimates for each of the 4 terms over the back half of iterations. F – I) Histograms of the estimates of the localization noise for each of the 4 terms over the back half of iterations. J – M) Histograms of the estimates of the weight fractions for each of the 4 terms over the back half of iterations. N – Q) Histograms of the estimates for the transition matrix elements giving the probability that a step in Term 1 is followed by a step in Term 1 on the next step (N), that a step in Term 1 transitions to a step in Term 2 (O), that a step in Term 1 transitions to a step in Term 3 (P), or that a step in Term 1 transitions to a step in Term 4 (Q). Black arrows in B – Q are the true simulation values (Table S1).

**Figure S2.**
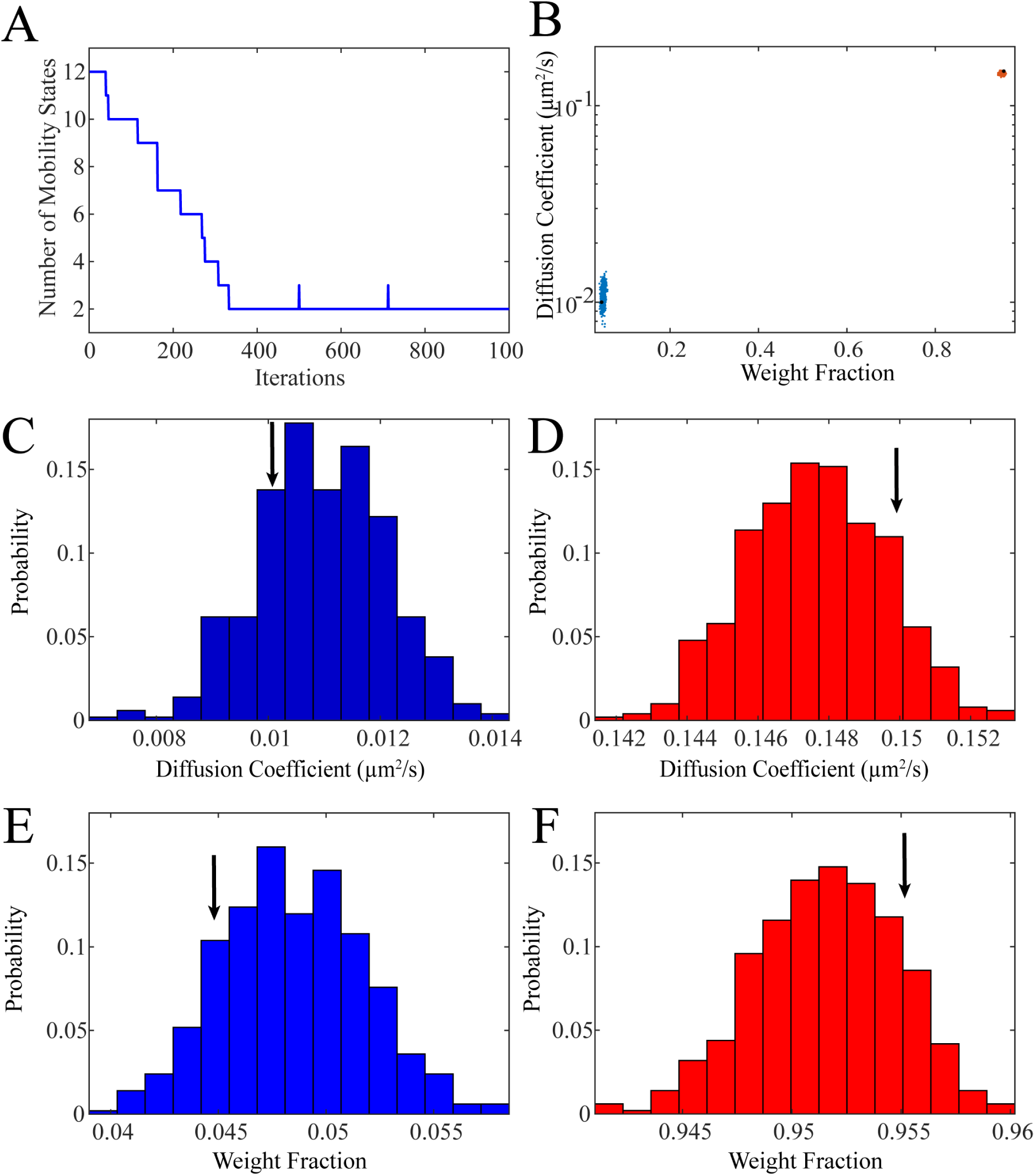
Full SMAUG analysis for the rare states simulation. A) Estimated mobility states over the course of the analysis run. SMAUG quickly converges to the correct value of *K* = 2, but continues to explore alternative hypotheses stochastically. B) Diffusion coefficient and weight fraction estimates for each saved iteration in the back half of the analysis run that also meets the *K* = 2 criterion. Black dots are the true simulation values. C – D) Histograms for the estimated diffusion coefficient values for the simulation. E – F) Weight fraction estimates. Black arrows are the true simulation values (Table S2).

**Figure S3.**
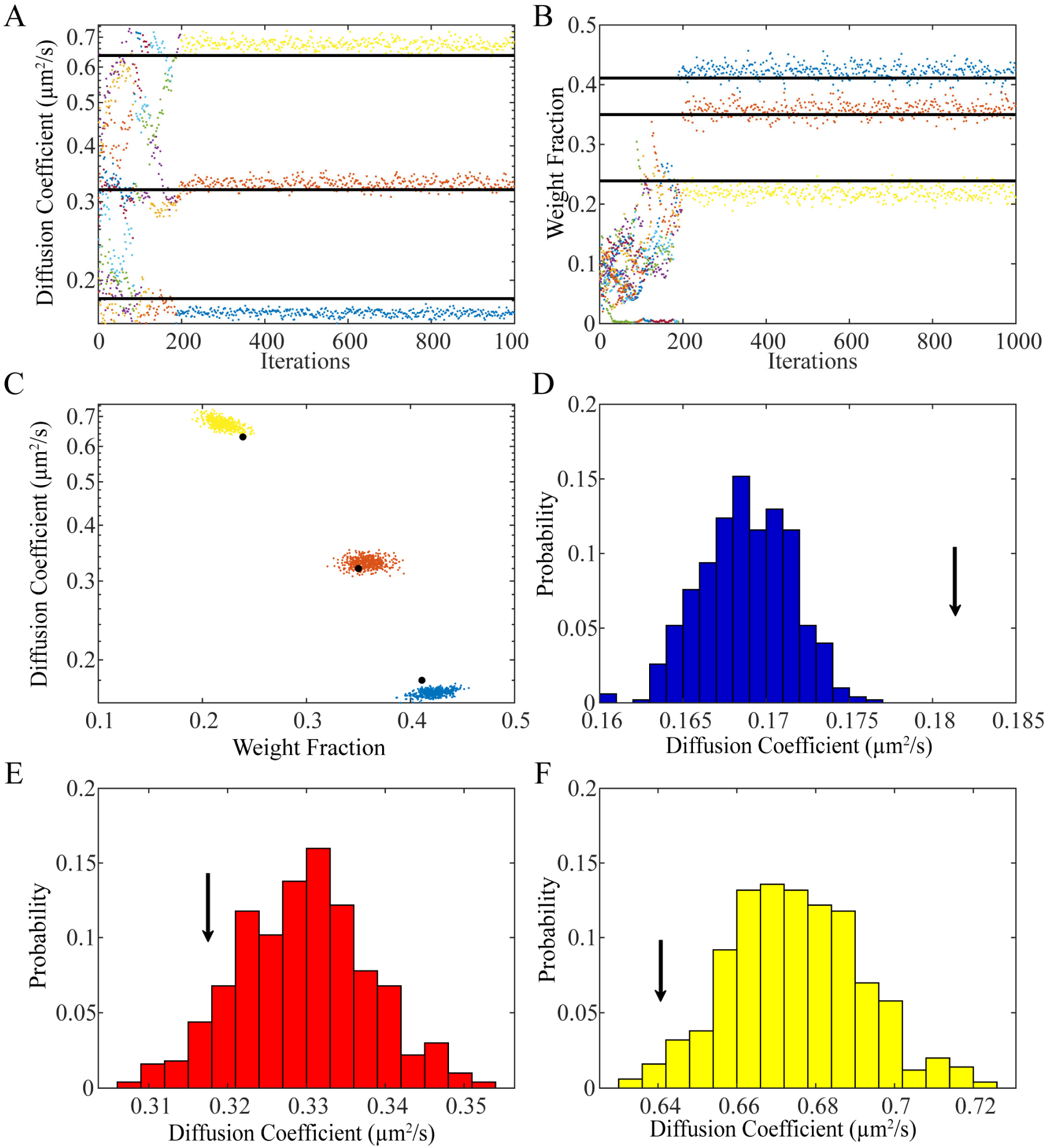
Full SMAUG analysis of the beads diffusing in glycerol *in vitro* experiments. A) Diffusion coefficient estimates for the analysis run. Black lines are the theoretical values for diffusion of beads in 50% glycerol. B) Weight fraction estimates for the analysis. Black lines are the true value of the number of steps from each size of bead. C) Diffusion coefficient and weight fraction estimates for each saved iteration in the back half of the analysis run that also meets the *K* = 3 criterion. Black dots correspond to the black lines in ‘A’ and ‘B’. D – F) histograms for the diffusion coefficient estimates. Black arrows represent the theoretical values (Table S3).

**Figure S4.**
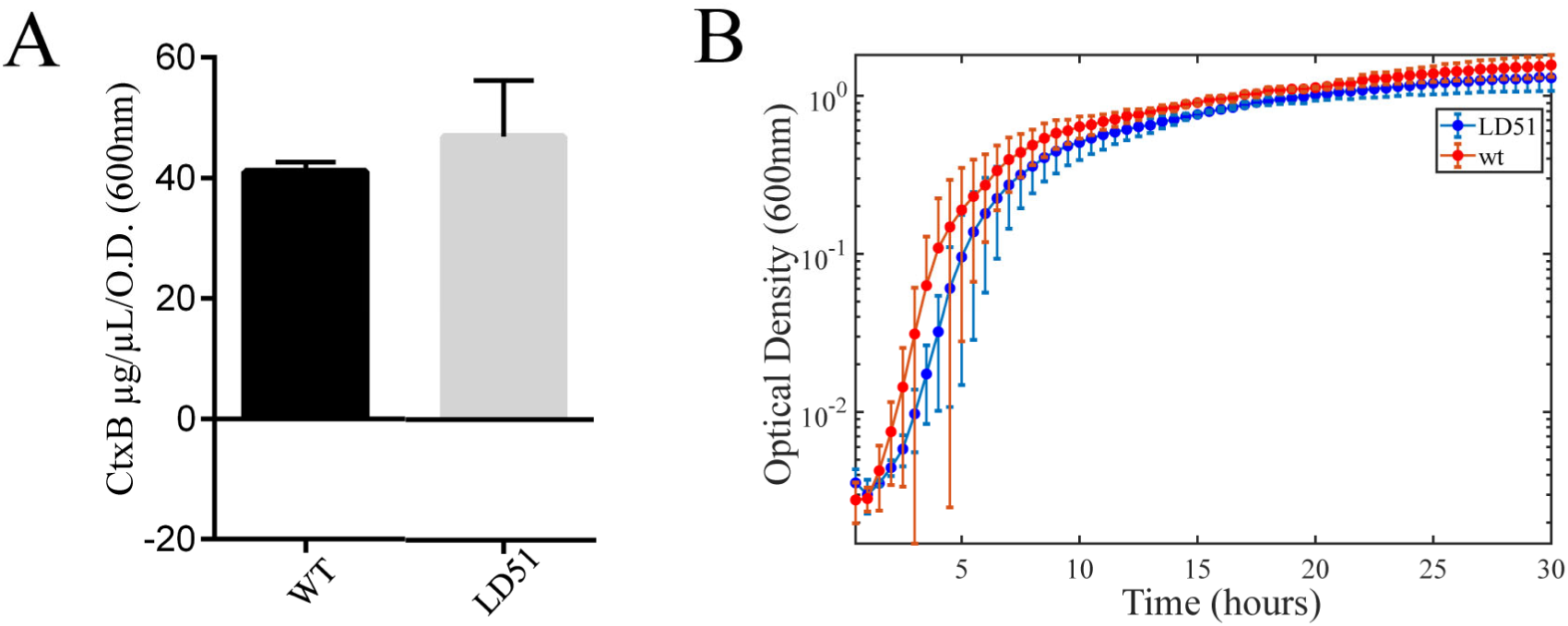
A) CtxB levels in culture supernatants after 24 h in LB media (pH 6.5, 30 ºC) show that LD51 expresses the same amount of CtxB protein as wild-type (WT) *V. cholerae* cells. B) Growth of WT and LD51 cells in LB (pH 6.5, 30 ºC). OD600 nm values are an average of three biological replicates.

**Figure S5.**
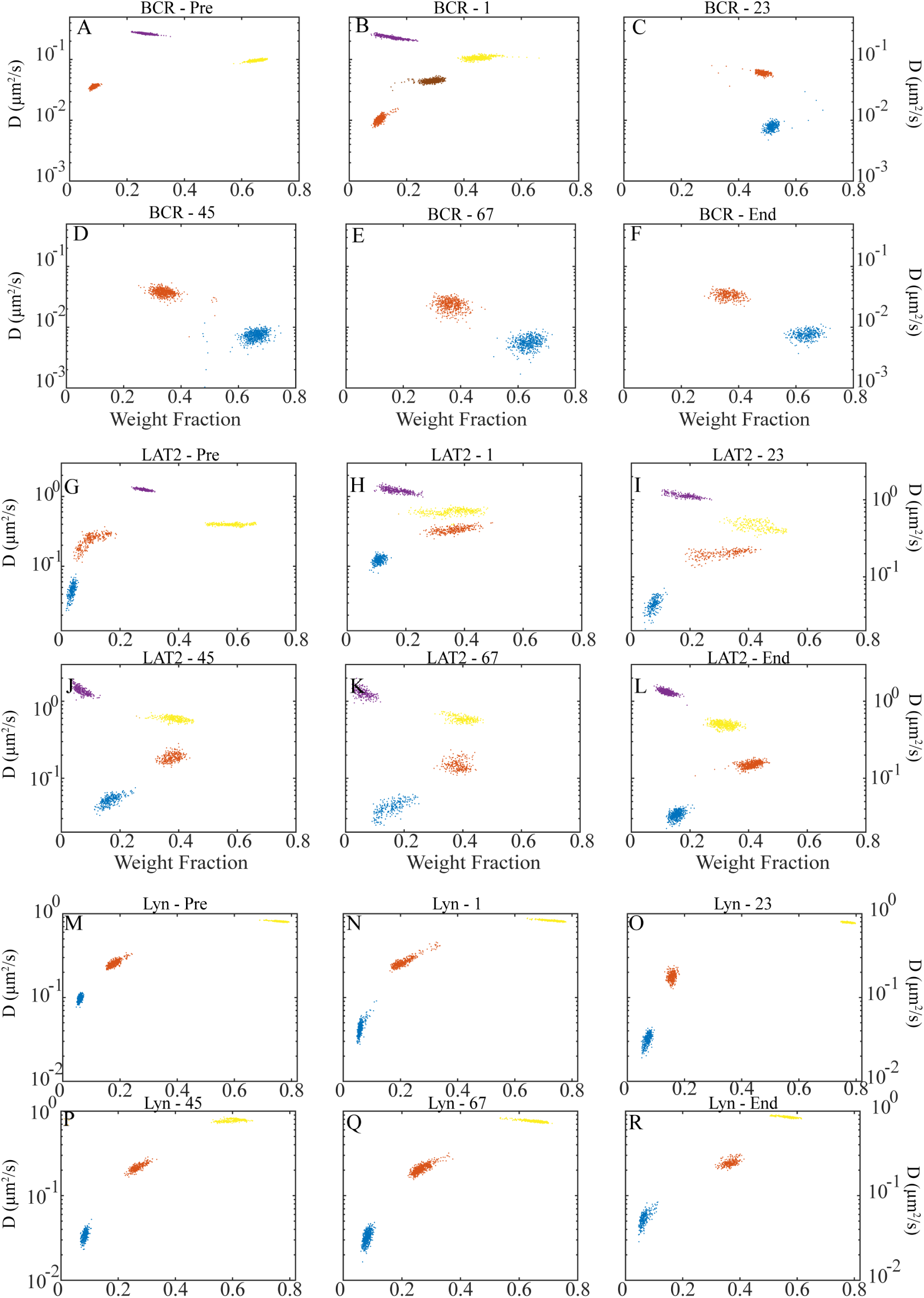
Full cluster analysis for the BCR, LAT2 and LYN molecules. A – F) SMAUG analysis of the diffusion coefficients and weight fractions for BCR-SiR. ‘A’ corresponds to the ‘Pre’ bar in Figure 4D, ‘B – E’ correspond to the middle bars, and ‘F’ corresponds to End. ‘A’ and ‘F’ are the same as in Figure 4C. G – L) SMAUG analysis of the diffusion coefficient and weight fraction for the LAT2. G corresponds to the ‘Pre’ bar in Figure 4E, ‘H – K’ correspond to the middle bars and L corresponds to End. M – R) SMAUG analysis of the diffusion coefficient and weight fraction for LYN. ‘M’ corresponds to the ‘Pre’ bar in Figure 4F, ‘N – Q’ correspond to the middle bars and ‘R’ corresponds to End.

**Figure S6.**
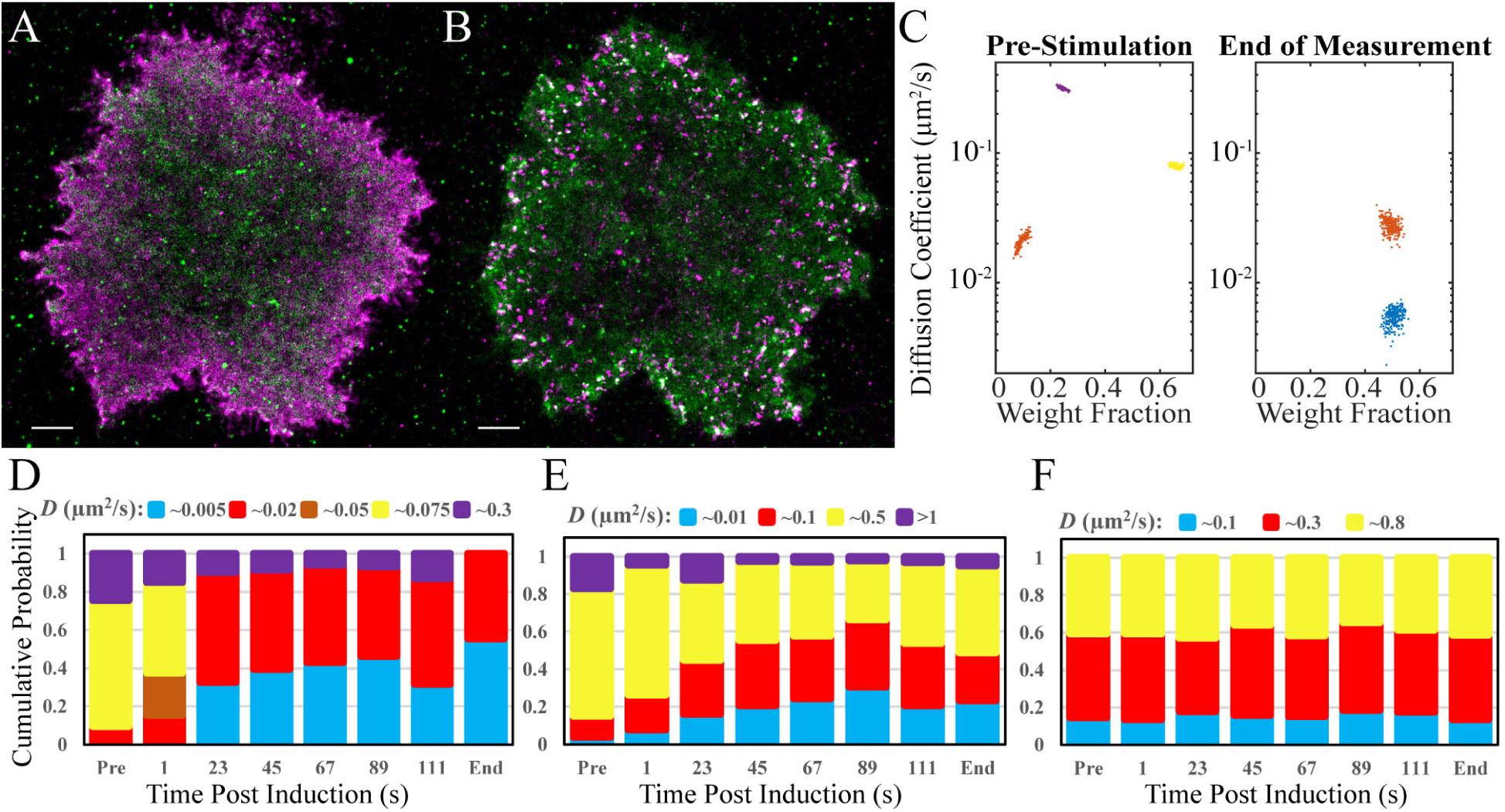
SMAUG analysis for single-molecule motion in a second B cell. A) Super-resolution reconstruction image of BCR-SiR (magenta) and LAT2-mEos3.2 (green) in a representative B cell pre-stimulation. Scale bar: 2 µm. B) Super-resolution image of the cell in ‘A’ 12.8 min post-stimulation. Scale bar: 2 µm. C) Diffusion coefficient and weight fraction estimates for BCR molecules pre-stimulation and at the end of the measurement. D) Bar graphs showing the mean weight fraction of each identified state as a function of time for the BCR dataset. The bars labeled “Pre” and “End” correspond to the data in ‘C’. All other bars are labeled with the time post-stimulation. Identified mobility states are states whose estimates overlap in diffusion coefficient and weight fraction. E) The bar graphs for the weight fractions of LAT2. F) The bar graphs for the weight fractions of Lyn. Analysis shows similar results to the cells used in Figure 4.

