## Supplemental - SMAUG Code for "SMAUG: Analyzing single-molecule tracks with nonparametric Bayesian statistics": README.txt

# SMAUG
Code needed for running the SMAUG analysis program.  


Single-Molecule Analysis by Unsupervised Gibbs sampling accurately and precisely determines estimates for several parameter values from single-molecule tracking data using a Bayesian statistical framework. 


Written by Josh Karslake at the University of Michigan.

## Installation

Download the entire zip folder and unzip. Change the working directory in Matlab to this folder. 

## Usage

SMAUG is an self-contained code suite for analysis of single-molecule tracking data. 

If you are using the output from the Biteen Lab SMALL-LABS (https://github.com/BiteenMatlab/SMALL-LABS), simply call the main function (‘SMAUG’) in the Matlab command window and then navigate through your folders to multi-select the "_fits" files, which contain the tracks.  

You can change the parameter values inside the code, such as integration time of the camera ('ImgIntTime') or max iterations by either by using paired inputs (SMAUG(‘ImgIntTime’,0.04)) or by changing the value itself inside the SMAUG function and saving. 

If you are using data files of your own creation, then some formatting will get the data into SMAUG-readable format. SMAUG needs 5 columns of data: 
-column 1: track number
-column 2: step number within that track
-column 3: a placeholder I used for some simulations and testing
-column 4: x position (in pixels)
-column 5: y position (in pixels)

For instance, in the sample data file 'SMAUG_TestTracks.mat', track 1 consists of 11 localizations. The first localization is at the position (x,y) = (0.3775 px, -0.1218 px).

NOTE: the x and y positions are in units of PIXELS by default and get scaled by ImgNPP (image nanometers per pixel) later, so if your localizations are already in nanometers, change Params.ImgNPP to 1 with the command (SMAUG(‘ImgNPP’,1)).

## Contributing

Please inform us before making any changes, then follow the directions below: 
1. Fork it!
2. Create your feature branch: `git checkout -b my-new-feature`
3. Commit your changes: `git commit -am 'Add some feature'`
4. Push to the branch: `git push origin my-new-feature`
5. Submit a pull request :D

## Credits
The development of this code is greatly indebted to the work of David J. Rowland and also to Keegan E Hines. The MakeSteps function was used from David’s CPDGlobal function (https://github.com/BiteenMatlab/SingleMoleculeDataAnalysis - Rowland and Biteen, Chemical Physics Letters, 674 173-178 (2017)), and parts of the the latent variable calculation, hyper-parameter calculations, and some skeleton organization were taken from/inspired by Hines, Biophysical Journal, 108 2103-2113 (2015). 


## License

                      GNU GENERAL PUBLIC LICENSE
                       Version 3, 29 June 2007

  See LICENSE.txt
