## Supplemental - SMAUG Code for "SMAUG: Analyzing single-molecule tracks with nonparametric Bayesian statistics": SMAUG.m

```
function [out]=SMAUG(varargin)

%% Parameters
disp('Parameters.......set')%this is an inside joke
disp('LOADING')

%imaging parameters
Params.ImgIntTime=0.04;     % frame integration time in seconds
Params.ImgNPP=49;           % Nanometeres per pixel .082; %60x value  .049 100x value 160 sarah

%analysis parameters
Params.MinTrLength=4;     %min number of localizations for a track to be considered
Params.IntL=10;           %initial number of states
Params.MaxIter=10000;     %max iterations to consider after which SMAUG will hard stop
% Params.IterStop=5000;    %someties I like to have it stop and check things as they go. set to 0 if you want to not do this
Params.IterSaveFreq=10;  %#iterations between saves to the output file, gotta be nonzero
Params.BurnIn=500;       %burn-in  none of the calculations on the chains themselves will kick in until after the burn-in
% Params.Resort=1;          % do you want to occasionally scramble the track order.
% Params.ResortFreq=10;     %how often do you want to scramble it up,
Params.Bootstrap=0;       %bootstrapping of the datasets by resample with replacement
Params.ResampleHypers=0;  % toggle to resample the hyper parameters using the hyper-hyper parameters
Params.ResampHypersFreq=5;% set to how many iterations before the hypers are resampled

% toggles for various outputs
Params.YesPlot=1;         %do you want to plot the output when its done
Params.YesMatfile=1;      % do you want to save to a matfile the analysis?
Params.SaveWorkSpace=0;   %if 1 will save the whole workspace, bigger files but maybe you want to save the whole thing
Params.MatfileName='name of analysis out file'; %if you want to output a file got to give it a name

%wheres the data to be analyzed 
Params.TrackFileLoc=[];   %location of a folder holding many files
Params.TrackFile=[];         %leave empty to select data


%hypervariables for the code. most are set to be flat. if desired to have
%an adaptive hyperparameter toggle Params.ResampleHypers to 1 above
Hypers.a0=1;
Hypers.g=.1;
Hypers.hyp_aa=1;
Hypers.hyp_ab=1;
Hypers.hyp_ga=1;
Hypers.hyp_gb=1;
Hypers.NormA=.01;
Hypers.NormB=.01;
Hypers.NormK=1;
Hypers.NormMu=0;
Hypers.CorrA=1;
Hypers.CorrB=1;
Hypers.CorrMu=0;%-10;
Hypers.CorrK=1;%500;
Hypers.Num=25;

%% you can change the Params file from the command line if you want using paired inputs
fNames=fieldnames(Params);
if nargin>1&&rem(nargin,2)==0
    for ii=1:2:nargin-1
        whichField = strcmp(fNames,varargin{ii});
        if all(~whichField)
            warning(['Check spelling. ', ...
                'Parameter change may have not occurred'])
        end
        eval([fNames{whichField} ' = varargin{ii+1};'])
        eval(['Params.' fNames{whichField} ' = ' fNames{whichField},';'])
    end
elseif nargin>1
    warning('use paired inputs')
    return
end
%% Make the dataset of steps from the tracks
[Steps,CorrSteps,dnames]=SMAUG_MakeSteps(Params);

%% Set up the iterations
out.Sample={};
[Sample]=SMAUG_Setup(Steps,CorrSteps,Params,Hypers); %first iteration and some other bits
fprintf('Iteration: %d: L = %d \n',Sample.Iter, Sample.L);

while Sample.Iter<Params.MaxIter %the bulk of the calcs
    
    u=zeros(1,length(Sample.l));
    u(1)=rand*Sample.TM(1,Sample.l(1)); %allowable states for u
    for ii=2:length(Sample.l)
        u(ii)=rand*Sample.TM(Sample.l(ii-1),Sample.l(ii));
    end
    Sample.U=u;
    
    while max(Sample.TM(:,end))>min(u) %add term(s) as needed
        [Sample]=SMAUG_AddTerms(Sample,Hypers);
    end
    Sample.L=size(Sample.TM,1);
    
    %forward filter, backward selector for the labels
    [Sample]=SMAUG_SampleHiddenStates(Sample,Steps,Params);
    
    
    %resample the params and hypers based on the new assignments
    [Sample]=SMAUG_Hypers(Sample,Params,Hypers);
    [Sample]=SMAUG_CalcParams(Steps,CorrSteps,Sample,Hypers);
    
    
    %clean up the states space
    [Sample]=SMAUG_CleanUp(Sample,Params);
    
    
    %saving (convergence testing added in later)
    if rem(Sample.Iter,Params.IterSaveFreq)==0
        [out,Sample]=SMAUG_Save(Sample,Steps,Params,out,dnames);
    end
    
    if rem((Sample.Iter/50),1)==0
        fprintf('Iteration: %d: L = %d ; D=\n ',Sample.Iter, Sample.L);
        disp(sort(out.Dvals{Sample.isave-1}'))
    end
    
    %iterstop things?
%     if rem((Sample.Iter/4500),1)==0
%         keyboard
%     end
    Sample.Iter=Sample.Iter+1;
end

%% output stuff and plotting
if Params.YesMatfile
    if Params.SaveWorkSpace==1 %whole thing
        save(Params.MatfileName);
    else   % or just the most important bits
        mm=matfile(Params.MatfileName,'Writable',true);
        mm.out=out;
    end
end

%making various figures of the output 
if Params.YesPlot
    SMAUG_Plot(Sample,out)
end

end  
```
