## Supplemental - SMAUG Code for "SMAUG: Analyzing single-molecule tracks with nonparametric Bayesian statistics": SMAUG_AddTerms.m

```
function [Sample]=SMAUG_AddTerms(Sample,Hypers)
% a function to add a term to the SMAUG analysis by drawing a random value
% from P0 and then breaking the beta stick again to assign that parameter
% some weight

%add another value to the end of the current Sample parameters
AddTerm=length(Sample.Beta);
Sample.TM(AddTerm,:)=drchrnd(Sample.Alpha*Sample.Beta,1);
Sample.Sigma(AddTerm)=1/gamrnd(Hypers.hyp_ga,Hypers.hyp_gb);
Sample.Mu(AddTerm)=randn*sqrt(Sample.Sigma(AddTerm))+Hypers.NormMu;

%Break the stick again
OldB=Sample.Beta(AddTerm);
NewB=betarnd(1,Sample.Gamma);
Sample.Beta(AddTerm)=NewB*OldB;
Sample.Beta(AddTerm+1)=(1-NewB)*OldB;

OldTM=Sample.TM(:,end);
A=repmat(Sample.Alpha*Sample.Beta(end-1),AddTerm,1);
B=Sample.Alpha*(1-sum(Sample.Beta(1:end-1)));
NewTM=betarnd(A,B);
if isnan(sum(NewTM))
    NewTM=binornd(1,A./(A+B));
end
Sample.TM(:,AddTerm)=NewTM.*OldTM;
Sample.TM(:,AddTerm+1)=(1-NewTM).*OldTM;
end
```
