## Supplemental - SMAUG Code for "SMAUG: Analyzing single-molecule tracks with nonparametric Bayesian statistics": SMAUG_CalcParams.m

```
function [Sample]=SMAUG_CalcParams(Steps,CorrSteps,Sample,Hypers)
%SMAUG_SampleNormal calculates the parameter values for a normal inverse
%gamma distribution using the data assignments 

Sample.Sigma=zeros(max(Sample.l),1);
Sample.Mu=zeros(max(Sample.l),1);
Sample.CSigma=zeros(max(Sample.l),1);
Sample.CMu=zeros(max(Sample.l),1);
%calculate parameters for the normal corresponding to step size
%distribution
for ii=1:max(Sample.l) 
    Data=Steps(Sample.l==ii,1:2);
    Data=cat(1,Data(:,1),Data(:,2));
    N=size(Data,1);
    if N==0
        Sample.Sigma(ii)=1/gamrnd(Hypers.hyp_ga,Hypers.hyp_gb);
        Sample.Mu(ii)=randn(1)*Sample.Sigma(ii)+Hypers.NormMu;
    else
        ML=nanmean(Data);
        Ahat=Hypers.NormA+N/2;
        Bhat=Hypers.NormB+.5*nansum((Data-ML).^2)+...
            ((N*Hypers.NormK)/(2*(N+Hypers.NormK))*(ML-Hypers.NormMu)^2);
        Sample.Sigma(ii)=sqrt(1/gamrnd(Ahat,1/Bhat));
        Tau=1/(Sample.Sigma(ii)^2);
        Khat=(N*Tau+Hypers.NormK*Tau)^-1;
        Mhat=((N*Tau/(N*Tau+Hypers.NormK*Tau))*ML)+...
            ((Hypers.NormK*Tau/(N*Tau+Hypers.NormK*Tau))*Hypers.NormMu);
        Sample.Mu(ii)=randn(1)*sqrt(Khat)+Mhat;
    end
end
%calculate the parameters corresponding to the correlated step sizes
%distribution
for ii=1:max(Sample.l)
    CData=CorrSteps(Sample.l==ii,1:2);
    CData=cat(1,CData(:,1),CData(:,2));
    N=size(CData,1);
    if N==0
        Sample.CSigma(ii)=1/gamrnd(Hypers.hyp_ga,Hypers.hyp_gb);
        Sample.CMu(ii)=randn(1)*Sample.CSigma(ii)+Hypers.CorrMu;
    else
        CML=nanmean(CData);
        Ahat=Hypers.CorrA+N/2;
        Bhat=Hypers.CorrB+.5*nansum((CData-CML).^2)+...
            ((N*Hypers.CorrK)/(2*(N+Hypers.CorrK))*(CML-Hypers.CorrMu)^2);
        Sample.CSigma(ii)=sqrt(1/gamrnd(Ahat,1/Bhat));
        Tau=1/(Sample.CSigma(ii)^2);
        Khat=(N*Tau+Hypers.CorrK*Tau)^-1;
        Mhat=((N*Tau/(N*Tau+Hypers.CorrK*Tau))*CML)+...
            ((Hypers.CorrK*Tau/(N*Tau+Hypers.CorrK*Tau))*Hypers.CorrMu);
        Sample.CMu(ii)=randn(1)*sqrt(Khat)+Mhat;
    end
end
%used to calculate allowable transitions 
T2=padarray(Sample.TM,[1,1],0,'post');
for jj=1:length(Sample.Beta)
    AllowedTrans(jj,:)=drchrnd((T2(jj,:)+(Sample.Alpha*Sample.Beta)),1);
end
AllowedTrans(end,:)=[];
Sample.TM=AllowedTrans;
Sample.L=size(AllowedTrans,1);

end
```
