## Supplemental - SMAUG Code for "SMAUG: Analyzing single-molecule tracks with nonparametric Bayesian statistics": SMAUG_CleanUp.m

```
function [Sample]=SMAUG_CleanUp(Sample,Params)
% this function will remove terms from the state space that are no longer
% needed

if Sample.Iter>Params.BurnIn
    Dvals=(Sample.Sigma.^2+2*Sample.CMu)/2/Params.ImgIntTime;
    if any(diff(sort(Dvals))<25)
        [~,i2]=sort(Dvals);
        Rt=find(diff(sort(Dvals))<25);
        for ii=1:length(Rt)
            Sample.l(Sample.l==Rt(ii))=i2(find(i2==Rt(ii)))+1;
        end
    end    
    if any(Dvals<0)&&Sample.L>2
        [~,i2]=sort(Dvals);
        Sample.l(Sample.l==i2(1))=i2(2);
    end
end

BadTerm=sort(setdiff(1:Sample.L,unique(Sample.l)));
for zz=length(BadTerm):-1:1
    Sample.Beta(end)=Sample.Beta(end)+Sample.Beta(BadTerm(zz));
    Sample.Beta(BadTerm(zz))=[];
    Sample.TM(:,BadTerm(zz))=[];
    Sample.TM(BadTerm(zz),:)=[];
    Sample.Mu(BadTerm(zz))=[];
    Sample.Sigma(BadTerm(zz))=[];
    Sample.CSigma(BadTerm(zz))=[];
    Sample.CMu(BadTerm(zz))=[];
    Sample.Pi(BadTerm(zz))=[];
    Sample.l(Sample.l>BadTerm(zz))=Sample.l(Sample.l>BadTerm(zz))-1;
end
Sample.L=size(Sample.TM,1); 
end
```
