## Supplemental - SMAUG Code for "SMAUG: Analyzing single-molecule tracks with nonparametric Bayesian statistics": SMAUG_Hypers.m

```
function [Sample]=SMAUG_Hypers(Sample,Params,Hypers)
% SMAUG_Hypers calculates the hyper parameters adapted from code from Hines
% 2016

T=zeros(max(Sample.l));
T(1,Sample.l(1))=1;
for tt=2:size(Sample.l,2)
    T(Sample.l(tt-1),Sample.l(tt))=T(Sample.l(tt-1),Sample.l(tt)) + 1;
end
Dishes = zeros(max(Sample.l));
for ii=1:max(Sample.l)
    for jj=1:max(Sample.l)
        if T(ii,jj)==0
            Dishes(ii,jj)=0;
        else
            for kk=1:T(ii,jj)
                Dishes(ii,jj)=Dishes(ii,jj)+(rand()<(Sample.Alpha*Sample.Beta(jj))/(Sample.Alpha*Sample.Beta(jj)+kk-1));
            end
        end
    end
end
Sample.Beta=drchrnd([sum(Dishes,1),Sample.Gamma],1);
Sample.TM=T;


if Params.ResampleHypers==1 %resample a0 and gamma from the hyper-hypers
    if rem(Sample.Iter,Params.ResampHypersFreq)==0
        D = sum(sum(Dishes));
        k = length(Sample.Beta);
        for iter = 1:Hypers.Num
            b1=sum(log(betarnd(Sample.Alpha+1,sum(T,2))));
            p1=sum(T,2)/Sample.Alpha; p1=p1./(p1+1);
            s1=sum(binornd(1,p1));
            Sample.Alpha=gamrnd(Hypers.hyp_aa+D-s1,1.0/(Hypers.hyp_ab-b1));
            Mu=betarnd(Sample.Gamma+1,D);
            if rand() < 1/(1+(D*(Hypers.hyp_gb-log(Mu)))/(Hypers.hyp_ga+k-1))
                Sample.Gamma=gamrnd(Hypers.hyp_ga+k,1.0/(Hypers.hyp_gb-log(Mu)));
            else
                Sample.Gamma=gamrnd(Hypers.hyp_ga+k-1,1.0/(Hypers.hyp_gb-log(Mu)));
            end
        end        
    end   
end
end
```
