## Supplemental - SMAUG Code for "SMAUG: Analyzing single-molecule tracks with nonparametric Bayesian statistics": SMAUG_MakeSteps.m

```
function [Steps,CorrSteps,dnames]=SMAUG_MakeSteps(Params)
%Makes matrix of step sizes for use in the SMAUG algorithm and keeps track
%of some other informatino for later

if isempty(Params.TrackFile)&&isempty(Params.TrackFileLoc)
    [TrFileName,trFileLoc,idf]=uigetfile('*.mat;*.dat;*.txt','Select analysis files.',...
        'MultiSelect','on');
    if ~idf
        display('no files chosen.')
        return
    end
    if ~iscell(TrFileName)
        TrFileName={TrFileName};
    end
    TrFileName = cellfun(@(x)[trFileLoc, x],TrFileName,'uniformoutput',false);
    [dlocs,dnames,dexts]=cellfun(@fileparts,TrFileName,'uniformoutput',false);
    %     if strcmp(dexts,'.mat')
    %         m=matfile(TrFileName{1},'Writable',true);  %this is a failsafe to make sure the int time is correct
    %         intTime=m.paramsTr;
    %         Params.ImgIntTime=intTime;
    %     end
elseif isempty(Params.TrackFile)&&~isempty(Params.TrackFileLoc) %open all the analysis files in a certain location
    cd(Params.TrackFileLoc)
    TrList=cat(1,dir('*_analysis*'),dir('*_fits*'));
    TrFileName={TrList(:).name};
    TrFileName = cellfun(@(x)[Params.TrackFileLoc,filesep, x],TrFileName,'uniformoutput',false);
    [dlocs,dnames,dexts]=cellfun(@fileparts,TrFileName,'uniformoutput',false); 
else
    TrFileName={'manual input'};
end


counter=0;
for kk=1:numel(TrFileName)
    if ~strcmp(TrFileName{kk},'manual input')
        if strcmp(dexts{kk},'.mat')==1&&strcmp(dnames{kk}(end-7:end),'analysis')%legacy masterfit3 output format
            m1=matfile([TrFileName{kk}]);
            try
                trfile=m1.trackfile;
            catch
                disp([TrFileName{kk} ' does not include tracking data. Skipping'])
                continue
            end
        elseif strcmp(dexts{kk},'.dat')==1 %even older Yi's code
            fid=fopen(TrFileName{kk},'rt');
            trfile=textscan(fid,'%f%f%f%f%f%f%f%f%f%f%f%f%f%f%f%f','HeaderLines',1,'CollectOutput',1);
            trfile=trfile{1};
            fclose(fid);
        elseif strcmp(dexts{kk},'.txt')==1 %sarahs stuff
            fID=fopen(TrFileName{kk},'r');
            fspec='%f %f %f %f %f';
            fsz=[5,Inf];
            tracks=fscanf(fID,fspec,fsz);
            trfile=tracks';
        elseif strcmp(dexts{kk},'.mat')==1&&strcmp(dnames{kk}(end-3:end),'fits') %SMALL-labs code
            load(TrFileName{kk},'tracks')
            trfile=tracks(:,[4,1,5,2,3]);
        elseif strcmp(dexts{kk},'.mat') %the simulations test code
            load(TrFileName{kk},'trfile')
        end
    else
        trfile=Params.TrackFile;
    end
    
    
    % loop through tracks
    for ii=unique(trfile(:,1))'
        % select current track from the array of all tracks
        trackii=trfile(trfile(:,1)==ii,:);
        if size(trackii,1)<Params.MinTrLength
            continue
        end
        counter=counter+1;
        % fill in the time holes with nans. these two lines are genius!
        % all hail the great dave
        fixedTrack=nan(max(trackii(:,2)),size(trackii,2));
        fixedTrack(trackii(:,2),:)=trackii;
        fixedTrack(:,[4,5])=fixedTrack(:,[4,5]).*Params.ImgNPP; %scale to nanometers from pixels
        % remove leading nans
        fixedTrack(1:find(all(isnan(fixedTrack),2)==0,1,'first')-1,:)=[];
        % overlapping frame pairs
        indvec1=2:size(fixedTrack,1);
        indvec2=1:size(fixedTrack,1)-1;
        
        % nansum because there are nans as placeholders
        TotSteps{counter,1}(:,1)=nansum((fixedTrack(indvec1,4)-...
            fixedTrack(indvec2,4)),2);
        TotSteps{counter,1}(:,2)=nansum((fixedTrack(indvec1,5)-... %these first 2 columns are steps
            fixedTrack(indvec2,5)),2);
        TotSteps{counter,1}(:,3)=diff(fixedTrack(:,2)); %this column shows which steps are adjacent in time
        TotSteps{counter,1}(:,4)=fixedTrack(1:end-1,3);
        TotSteps{counter,1}(:,5)=ones(size(fixedTrack(1:end-1,1))).*kk; %label which movie these tracks came from
        TotSteps{counter,1}(:,6)=ones(size(fixedTrack(1:end-1,1))).*ii; %label which track fom that movie these steps came from
    end
end


if Params.Bootstrap
    Steps=TotSteps(randsample(size(TotSteps,1),size(TotSteps,1),1),1);
else
    Steps=TotSteps(:,1);
end
Steps=Steps(~cellfun('isempty',Steps));

for ii=1:length(Steps)
    Steps{ii}=[Steps{ii}(:,1:2),ones(size(Steps{ii},1),1)*ii,Steps{ii}(:,3),Steps{ii}(:,5:6)];
end
Steps=cat(1,Steps{:});
Steps=Steps(Steps(:,1)~=0,:);Steps=Steps(Steps(:,2)~=0,:);
for ii=1:size(Steps,1)-1
    if Steps(ii,3)==Steps(ii+1,3)&&Steps(ii,4)==1
        CorrSteps(ii,1)=Steps(ii,1)*Steps(ii+1,1);
        CorrSteps(ii,2)=Steps(ii,2)*Steps(ii+1,2);
    else
        CorrSteps(ii,:)=nan;
    end
end
CorrSteps(end+1,:)=nan;
end
```
