## Supplemental - SMAUG Code for "SMAUG: Analyzing single-molecule tracks with nonparametric Bayesian statistics": SMAUG_Plot.m

```
function [out]=SMAUG_Plot(Sample,out)
%% plots some plots of the output structure


c=colormap('lines');

%plot the # of states per iteration
figure
plot(out.L,'b','linewidth',2)
title('Mobility Sates')
ylabel('Number of Sates')
xlabel('Iteration')

%scatter plot of Dvals vs iteration
figure
for ii=1:2:Sample.isave-1 %every other to save some time 
    Dsort=sort(out.Dvals{ii});
    for jj=1:length(Dsort)
        scatter(ii,Dsort(jj),7,c(jj,:),'filled')
        hold on
    end
end
set(gca,'yscale','log')
axis tight
title('Diffusion Value Estimates vs Iteration')
ylabel('Diffusion Coeffecient, \mum^2/s')
xlabel('Iterations')

%scatter plot of Dvals vs weight fraction for the most probable model
figure
for mm=round(Sample.isave/2):Sample.isave-1
    if out.L(mm)==mode(out.L(round(Sample.isave/2):end))
        [s2(mm,:),i2]=sort(out.Dvals{mm});
        w2(mm,:)=out.Pi{mm}(i2)/sum(out.Pi{mm});
    end
end
s2(s2(:,1)==0,:)=[];
w2(w2(:,1)==0,:)=[];
w3=mean(w2,1);
s3=mean(s2,1);
figure
hold on
for jj=1:mode(out.L(round(Sample.isave/2):end))
    scatter(w2(:,jj),s2(:,jj),7,c(jj,:),'filled')
end
set(gca,'yscale','log')
axis tight
titlestr=sprintf('Diffusion vs weight fraction for L = %d', mode(out.L));
title(titlestr)
ylabel('Diffusion Coeffecient, \mum^2/s')
xlabel('Weight Fraction')

end
```
