## Supplemental - SMAUG Code for "SMAUG: Analyzing single-molecule tracks with nonparametric Bayesian statistics": SMAUG_SampleHiddenStates.m

```
function [Sample]=SMAUG_SampleHiddenStates(Sample,Steps,Params)
%this function preforms a forward filter, backwards selector for each data
%point in the collection and then will assign that data point to a specific
%term

%likelihood calculations for each term using forward filter
labels=ones(1,length(Steps));
lkhood=zeros(Sample.L,length(Steps));
lkhood(:,1)=Sample.TM(1,1:Sample.L)>Sample.U(1);
for ii=1:Sample.L
    lkhood(ii,1)=(exp(-normlike([Sample.Mu(ii),Sample.Sigma(ii)],Steps(1,1:2))))*lkhood(ii,1);
end
lkhood(:,1)=lkhood(:,1)/sum(lkhood(:,1));

for jj=2:length(Steps)
    AllowedTrans=Sample.TM(1:Sample.L,1:Sample.L)>Sample.U(jj);
    lkhood(:,jj)=AllowedTrans'*lkhood(:,jj-1);
    for KK=1:Sample.L
        lkhood(KK,jj)=(exp(-normlike([Sample.Mu(KK),Sample.Sigma(KK)],Steps(jj,1:2))))*lkhood(KK,jj);
    end
    lkhood(:,jj)=lkhood(:,jj)/sum(lkhood(:,jj));
end

%backward selection
if sum(lkhood(:,end))~=0.0&&isfinite(sum(lkhood(:,end)))
    labels(end)=1+sum(rand()>cumsum(lkhood(:,end)));
    for t=length(Steps)-1:-1:1
        r = lkhood(:,t).*(Sample.TM(:,labels(t+1))>Sample.U(t+1));
        r = r./sum(r);
        labels(t)=find(mnrnd(1,r)==1);%1+sum(rand()>cumsum(r))
    end

    %use the label assignments to calculate the weights
    weights=zeros(1,Sample.L);
    for ii=unique(labels)
        weights(ii)=sum(labels==ii);
    end
    weights=drchrnd(weights,1/Sample.L);
    Sample.l=labels;
    Sample.LabelSaves(rem(Sample.Iter,Params.IterSaveFreq)+1,:)=labels;
    Sample.Pi=weights;
else
    fprintf('No path through the HMM. Reshuffling data to try again.\n');
    Sample.l=ceil(rand(1,length(Steps))*Sample.L);
    %make a counter so this doesnt happen too much??
end
end
```
