## Supplemental - SMAUG Code for "SMAUG: Analyzing single-molecule tracks with nonparametric Bayesian statistics": SMAUG_Save.m

```
function [out,Sample]=SMAUG_Save(Sample,Steps,Params,out,dnames)
%saves current values and things to the output structure


isave=Sample.isave;

out.L(isave)=Sample.L;
out.Labels{isave}=Sample.l;
% out.Sample{isave}=Sample;  files too big doing this, 
out.Dvals{isave}=(Sample.Sigma.^2+2*Sample.CMu)/2/Params.ImgIntTime/1e6; %converting to um^2/s
out.Pi{isave}=Sample.Pi'; %weight fractions
% out.TransMat{isave}=TMcalc(Steps(:,3),mode(Sample.LabelSaves,1),Sample.Alpha*Sample.Beta);
out.TransMat{isave}=TMcalc(Steps(:,3),mode(Sample.l,1),Sample.Alpha*Sample.Beta);
if Sample.Iter==1
    out.Params=Params;
    out.TrackIDs=Steps(:,5:6);
    out.MovNames=dnames;
    out.Params=Params;
end

Sample.isave=isave+1;

end
function [TM]=TMcalc(tracks,labels,H)
K = size(H,2);
TM = zeros(K);
N = zeros(K);
t=1;
for ii=1:max(unique(tracks))
    T=sum(tracks==ii);
    if T==0
        continue
    else
        t=t+1;
        for jj=2:T
            N(labels(t-1),labels(t))=N(labels(t-1),labels(t))+1;
            t=t+1;
        end
    end
end
for k=1:K
    TM(k, :)=drchrnd(N(k,:)+H,1);
end
TM(end,:)=[];
end
```
