## Supplemental - SMAUG Code for "SMAUG: Analyzing single-molecule tracks with nonparametric Bayesian statistics": SMAUG_Setup.m

```
function [Sample]=SMAUG_Setup(Steps,CorrSteps,Params,Hypers)
% SMAUGSetup sets up the structures needed in the rest of the code

Sample.l=ceil(rand(1,size(Steps,1))*Params.IntL); %randomly assigning steps into one of the IntL terms
weights=zeros(1,Params.IntL);
for ii=1:Params.IntL
    weights(ii)=sum(Sample.l==ii);
end
Sample.Pi=drchrnd(weights,1/Params.IntL);
Sample.Alpha=Hypers.a0;
Sample.Gamma=Hypers.g;
Sample.Beta=ones(1,Params.IntL)/Params.IntL;
Sample.Iter=1;
[Sample]=SMAUG_Hypers(Sample,Params,Hypers);
[Sample]=SMAUG_CalcParams(Steps,CorrSteps,Sample,Hypers);
Sample.isave=1;
Sample.LabelSaves=zeros(Params.IterSaveFreq,length(Steps));


%safety checks?


end
```
